# Molecular determinants of cardiac lymphatic dysfunction in a chronic pressure-overload model

**DOI:** 10.1101/2025.02.27.640360

**Authors:** C. Heron, T. Lemarcis, O. Laguerre, B. Bourgeois, C Thuilliez, C Valentin, A Dumesnil, M. Valet, D. Godefroy, D. Schapman, G. Riou, S. Candon, C. Derambure, A. Zernecke, C. Berard, H Dauchel, V. Tardif, E. Brakenhielm

## Abstract

Cardiac lymphatics have emerged as potential targets in cardiovascular diseases (CVDs). However, we recently reported that despite extensive lymphatic expansion during experimental cardiac pressure-overload, lymphatic drainage remained insufficient. To unravel the cellular and molecular mechanisms underlying lymphatic dysfunction in CVDs, we applied cardiac single-cell (sc) analyses in a murine heart failure model.

Transaortic constriction (TAC), in C57BL/6J and BALB/c mice, was used to model chronic pressure-overload-induced cardiac hypertrophy and heart failure, respectively. Cardiac lymphatic (LEC) and blood vascular (BECs) endothelial cells were analyzed by scRNAseq (10XGenomics). Lymphatic targets were validated by immunohistochemistry and wholemount-imaging, and *in vitro* using human LEC cultures.

We identified three distinct cardiac lymphatic subpopulations, capillary (*LEC1*), precollector (*LEC2*), and valvular (*LEC3*) clusters, and several BECs clusters, including venous BEC (*vBEC*). Chronic pressure-overload led to expansion of lymphatic capillaries and loss of valves in BALB/c, but not C75BL6/J. Analysis of differentially expressed genes (DEG) post-TAC revealed reduction only in BALB/c of lymphatic cell-junction components. In contrast, LEC expression of anchoring filaments, immune cell-adhesion molecules, and chemokines was preserved, or increased, indicating functional lymphatic-mediated immune cell uptake post-TAC. Interestingly, around 35% of DEGs identified in cardiac LECs post-TAC were similarly altered in interleukin (IL)-1β-stimulated human LECs.

In conclusion, loss of lymphatic valves and dysregulated lymphatic barrier properties may underly poor drainage capacity during pressure-overload, despite potent lymphangiogenesis and preserved LEC immune attraction. Further studies are needed to address how to restore lymphatic health to accelerate resolution of both inflammation and edema in CVDs.

## Introduction

Recent research has demonstrated that cardiac lymphatics remodel in CVDs in mice and men^1–3^. This includes not only structural changes, impacting lymphatic density and morphology, but also functional changes^4–6^, together determining cardiac lymphatic drainage capacity. Given that lymphatics are essential for cardiac homeostasis^7^, such development of cardiac lymphatic dysfunction, and/or insufficient lymphangiogenesis, in CVDs may accelerate cardiac inflammation, fibrosis, and adverse ventricular remodeling. Promisingly, therapeutic lymphangiogenesis has been reported to reduce cardiac dysfunction in models of myocardial infarction (MI) and chronic pressure-overload^4,8–14^. We recently demonstrated that pressure-overload-induced left ventricular (LV) dilation, following transaortic constriction (TAC), triggers massive endogenous cardiac lymphangiogenesis in BALB/c mice^15^. Intriguingly, the expanded lymphatic network failed to resolve chronic myocardial inflammation and edema, and the mice progressed to heart failure. We hypothesized that inflammatory mediators, including IL1β and IL6, produced in the heart during pressure-overload^15^, may modulate lymphangiogenesis and/or lymphatic function. Indeed, we recently demonstrated that IL1β plays a key role in induction of cardiac lymphangiogenesis post-TAC^16^. However, it remains unknown what cellular and molecular changes may contribute to poor lymphatic uptake and/or drainage during myocardial inflammation. To date, our knowledge of the molecular underpinnings of cardiac lymphatic (dys)function in CVDs is essentially limited to a few key players, including a cell junctional molecule (VE-Cadherin^6^) and select mediators of lymphatic-immune cell cross-talk (Lyve1^9^, CCL21^12,13,15,16^). Indeed, Lyve1 has been shown to be essential for cardiac immune cell exit through lymphatics post-MI, by interacting with hyaluronan-coated CD44-expressing immune cells to mediate leukocyte rolling and transmigration^9^. In contrast, pre-clinical investigations of lymphatics in other tissues, such as inflamed gut, skin, lymph nodes, and tumors, have revealed widespread transcriptional changes to lymphatic endothelial cells (LEC) during acute or chronic inflammation. Examples include alterations of capillary LEC cell junctions^9^; loss of lymphatic valves^17^; increased LEC expression of cell adhesion molecules (*Icam, Vcam*, *Selp*) and chemokines (*Ccl21, Cxcl12, Cx3cl1*)^18^; altered lymphatic production of bioactive lipids, e.g. sphingosine-1 phosphate (S1P)^19,20^; and finally increased lymphatic expression of immune-modulators, including major histocompatibility complex (MHC) for antigen-presentation, coupled to immune-suppressive co-signaling molecules, such as *Pdl1* (Programmed Death Ligand-1)^21^. Here, we set out to investigate the molecular changes that accompany cardiac lymphangiogenesis in the setting of inflammation induced by chronic pressure-overload, with the ultimate aim to uncover potential new targets, beyond lymphangiogenic growth factor therapy, to restore lymphatic drainage in CVDs. We also compared our data to published reports on lymphatic molecular profiles in other organs and disease settings to identify potential targets specific to cardiac lymphatics.

## Results

### Distinct blood vascular and lymphatic profiles in healthy mouse hearts

To define the transcriptome of healthy murine cardiac lymphatics, we used FACS to select cardiac CD45^-^/CD31^+^ endothelial cells (EC) from 20 sham-operated adult female BALB/c mice, further enriching the samples for Lyve1 and Podoplanin (Pdpn)-expressing LECs prior to scRNAseq (**Fig. 1a**). After quality controls (removing genes expressed by <10 cells; and removing cells expressing <500 genes, and >10% mitochondrial-encoded genes), we recovered 1,814 cardiac EC transcriptomes. These distributed (**Fig. 1b, c**) into three major clusters: *LEC* (402 transcriptomes), capillary/arterial blood vascular endothelial cells (*BEC*, 1,196 transcriptomes), and venous BECs (*vBEC*, 216 transcriptomes), as determined using Seurat graph–based unsupervised clustering and visualization by Uniform Manifold Approximation and Projection (UMAP). The EC clusters expressed classical vascular markers (*Pecam1, Kdr)* but not immune cell or smooth muscle cell markers (*Ptprc*, *Cd68, Acta2)* (**Suppl. Fig. S1a**). On average, in each cell, we detected 2,699 mRNA transcripts (reads), representing 1,341 unique genes (**Table S1**). The cell cycle phases of cardiac ECs were majoritarian G1 or G2/M, with slightly more BECs than LECs or vBECs in S phase in healthy adult mouse hearts (**Suppl. Fig. S1b, c**).

**Fig. 1.**
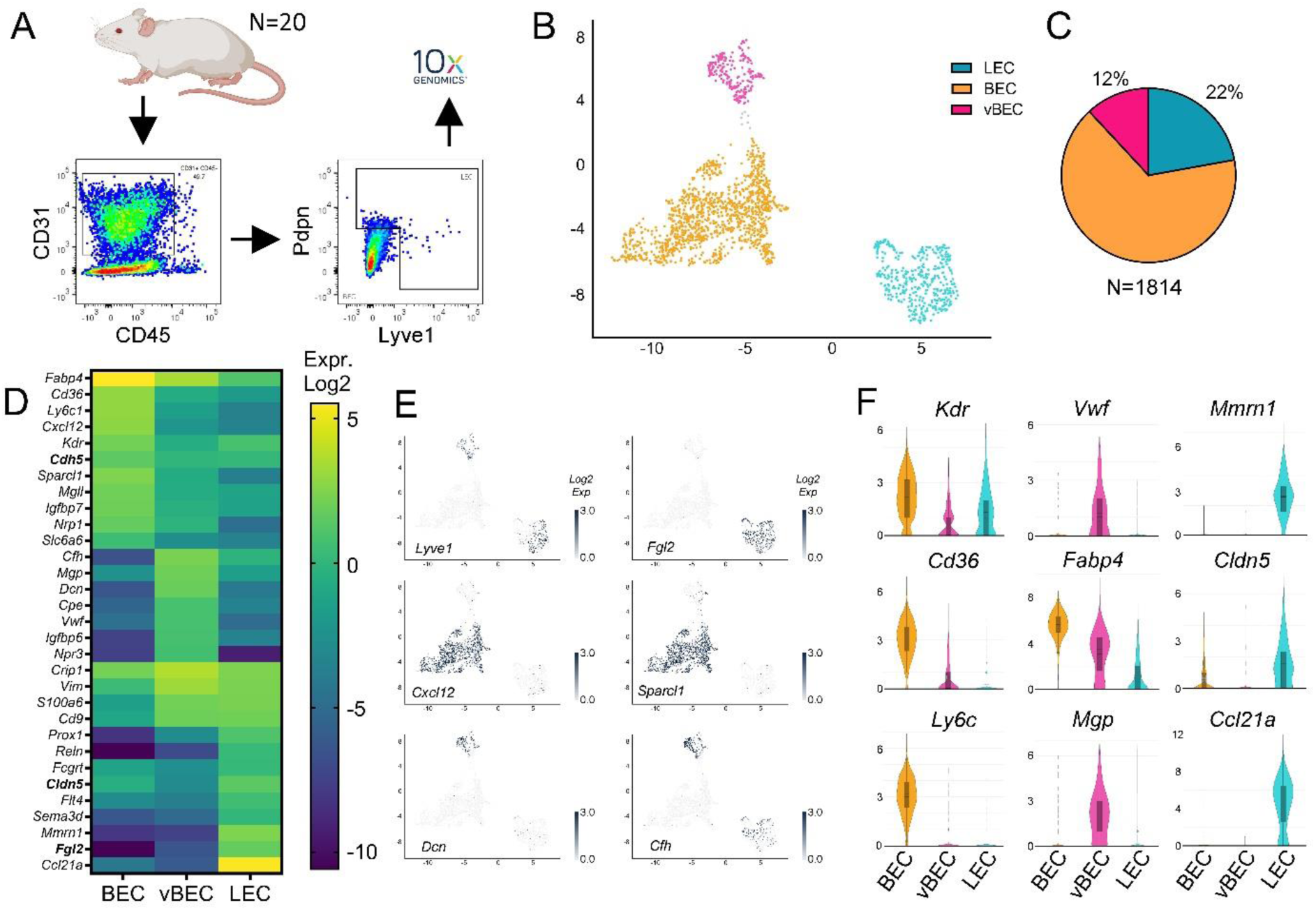
Distinguishing markers of healthy cardiac endothelial cell populations. Experimental overview: single cell preparations of cardiac cells from 20 healthy (sham-operated) mice were sorted by FACS to select global CD31^+^/CD45^neg^ cardiac endothelial cell (ECs) population and enrich samples for a rare subpopulation of ECs positive for LEC markers Lyve1 and Pdpn, before inclusion in 10x genomics pipeline (**a**). Unsupervised clustering [umap] of scRNAseq data, revealed three distinct cardiac vascular EC populations (*N=1814* transcriptomes): BEC (*orange*), LEC (*blue*), and vBEC (*pink*), representing 66%, 22%, and 12% of all ECs, respectively (**b, c**). Examples [heatmap; **d**], [umap, **e**], [violin plot, **f**] of expression profiles (gene expression levels shown as mean *Log2 normalized read counts*) selective for each cluster, including canonical lymphatic (*Ccl21, Mmrn1, Lyve1*, *Prox1, Flt4*); blood capillary and arteriolar (*Cxcl12, Nrp1, Kdr, Cd36)*; and venous (*Dcn*, *vWF*, *Mgp, Chf*) marker genes.

Comparing mean gene expression levels between cardiac EC clusters to identify population markers, we found 22, 45, and 35 genes distinguishing, respectively, LECs, BECs and vBECs (**Fig. 1d-f**; **Table S2**). BECs expressed higher levels, compared to the other clusters, of key blood vascular EC-selective genes, including *Cxcl12, Kdr* (VEGFR2)*, Nrp1* (Neuropilin-1)*, Aqp1* (Aquaporin-1), and the transcription factors *Id1* and *Ets1*. Cardiac BECs were further distinguished by elevated expression of lipolysis-related genes (**Fig. 1d, f**). Indeed, regulators of fatty acid (FA) uptake and cytosolic lipid trafficking, including *Lpl* (Lipoprotein Lipase), *Cd36,* and *Fabp4* (FA binding protein-4), were preferentially expressed in cardiac BECs (**Fig. 1f**, **Suppl. Fig. S2a**), while other key lipolysis-related genes, e.g. the mitochondrial lipid transporter *Cpt1a* (Carnitine palmitoyltransferase), were expressed at comparable levels in the cardiac EC clusters, as were several key glycolysis regulators (e.g. *Notch1*, *Foxo1*, *Hif1α*, *Slc2a1* (Glut1), *Slc2a8* (Glut8), *Hk1* (Hexokinase-1)) (**Suppl. Fig. S2a**). Our findings are in line with previous reports, comparing cardiac and brain ECs, revealing low levels of glucose transporters, but high levels of FA uptake genes in the cardiac vasculature^22^.

Cardiac vBECs were characterized by elevated expression of canonical venous markers, including *Dcn* (Decorin), *Vwf* (Von Willebrand Factor), and *Mgp* (Matrix Gla Protein), but also many other genes, such as *Npr3 (Natriuretic peptide receptor 3)* and *Cfh* (Complement factor H) (**Fig. 1d-f**; **Table S2**). vBECs were further distinguished by elevated expression of *H2az1* (MHC class I), compared to BECs and LECs, although *H2-D1* and *H2-K1* were the majoritarian MHC genes expressed in cardiac ECs (**Suppl. Fig. S2b**).

The cardiac LEC cluster was characterized by elevated expression of lymphatic markers, including *Ccl21a, Mmrn1* (Multimerin-1/Emilin4), *Flt4* (VEGFR3), and the transcription factors *Prox1, Nfat5/TonEBP,* and *Nfatc1* (**Fig. 1d-f, Table S2**). It also was distinguished by elevated expression of many other genes, including *Sema3d* (Semaphorin 3d), *Cd9* (tetraspanin family member), *Lbp* (Lipopolysaccharide-binding protein), *Fcgrt* (Fc gamma immunoglobulin receptor and transporter), and *Fgl2* (Fibrinogen-Like 2). The latter, expressed by the majority of cardiac LECs while barely detectable in BEC and vBEC clusters (**Fig. 1e**), encodes a secreted protein with immune-modulatory functions^23^. Of note, several of the identified non-canonical lymphatic genes expressed by cardiac LECs, including *Sema3d*, *Fgl2, Cd9,* and *Thy1* (CD90), have been previously detected in lymphatics in different organs in mice and humans^22,24,25^.

In the heart, maintenance of a tight blood vascular barrier represents a key function of ECs necessary for cardiac health, while lymphatic capillaries, in physiology, are permeable to allow fluid, macromolecule, and immune cell entry. Indeed, lymphatics are endowed with discontinuous button-junctions in capillaries, while precollectors display tight “zipper-type” junctions to promote efficient transport to lymph nodes^26^. In both blood vessels and lymphatics these vascular barriers are constituted by a combination of adherence (e.g. Cadherins, Catenins, Nectins) and tight junction (e.g. Claudins, Occludin, Esam, ZO) proteins (**Suppl. Fig. S3a, b**). We found in healthy hearts that these components displayed differential expression, with more *Cdh5* (VE-Cadherin) and *Jam2* (Junctional adhesion molecule B) in BECs, more *Cldn5* (Claudin-5) and *Nectin2* in LECs, and more *Cdh13* (T-Cadherin) in vBECs (**Fig. 1d, f**, **Suppl. Fig. S3a**). In contrast, gene expression levels of other junctional components, and of a major regulator of vascular barrier integrity, the S1P receptor *S1pr1,* were comparable between cardiac EC clusters. Reclustering analyses allowed investigations of BEC subpopulations (see **Table S3** for marker genes), revealing elevated *Cldn5* expression only in two clusters: BEC9 (**Suppl. Fig. S3c**), enriched for arteriolar markers (e.g. *Id1,* Notch-target gene *Hey1)*, and BEC6, enriched for angiogenic markers (e.g. *Aplnr2,* Apelin receptor 2).

### Identification of cardiac lymphatic subpopulations in healthy mice

Focusing next on cardiac LECs, reclustering analyses revealed three distinct subpopulations: LEC1 (65% of cells), LEC2 (22 % of cells), and LEC3 (13% of cells) (**Fig. 2a, b**). The majoritarian LEC1 cluster was distinguished by elevated expression of canonical lymphatic capillary markers, including *Mmrn1, Lyve1, Nrp2, Aqp1,* and *Flt4* (**Fig. 2c-e**, **Table S4**).

**Fig. 2.**
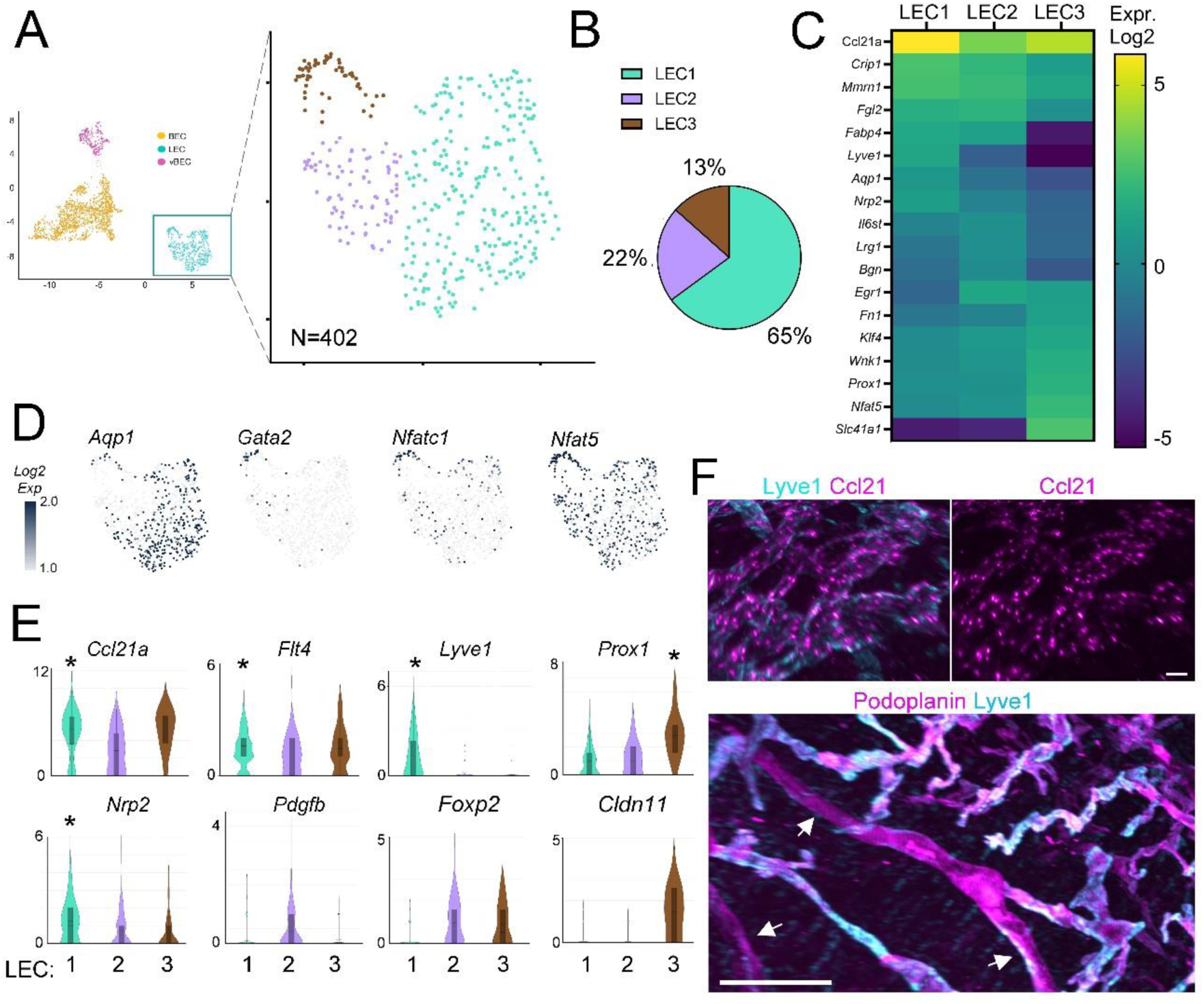
Identifying cardiac LEC subpopulations in healthy mouse hearts. Cardiac LECs (*N=402*) obtained from healthy sham-operated BALB/c mice (*n=20*) clustered (**a**) into three subpopulations: LEC1 (*cyan*), LEC2 (*purple*), and LEC3 (*brown*), representing 65%, 22% and 13% of all analysed LECs, respectively (**b**). Examples of mean expression levels of specific cluster markers (*Log2 normalized read counts*) [heatmap, **c**] [umap representation, **d**], [violin plot, **e**], including lymphatic capillary markers (*Ccl21, Mmrn1, Lyve1, Aqp1, Flt4, Nrp2*), precollector-enriched genes (*Pdgfb, Foxp2*), and lymphatic valve markers (*Gata2, Prox1, Nfat5, Cldn11)*. * denotes genes with significantly higher expression in a given cluster. Cardiac wholemount confocal imaging (**f**, *top panel*, x12, scalebar 30 µm) of CCL21-expressing lymphatic capillaries (Ccl21, *magenta*; Lyve1, *cyan*); and lightsheet imaging (**f**, *bottom panel*, x3.2, scalebar 300 µm) to illustrate lymphatic Lyve1^low^ precollector segments (Lyve1, *cyan*; Pdpn, *magenta*; white arrows point to LYVE1^low^ precollectors).

As mentioned previously, *Ccl21a* was highly expressed by all cardiac LECs, with the highest levels found in the capillary LEC1 cluster (**Fig. 2e**), similar as has been reported for lymphatics in other organs. Wholemount analyses further demonstrated steep lymphatic gradients of Ccl21 confined to the subepicardial regions of the heart, with a large proportion of the signal detected inside capillary LECs (**Fig. 2f**, **Suppl**. **video 1**).

The LEC2 subpopulation more frequently expressed *Pdgfb* (32% of cells) and *Foxp2* (55% of cells), as compared to LEC1 (8% and 13% of cells, respectively). The LEC2 cluster, which we identified as precollector-type, was also distinguished by higher expression of *Bgn* (Biglycan), *Cfh*, *Egr1* (Early Growth Response 1), and *Selenop* (encoding a secreted antioxidant selenoprotein), and lower expression of *Lyve1* and *Ccl21a*, as confirmed by confocal and lightsheet imaging (**Fig. 2c-f**, **Table S4**).

The small LEC3 cluster was characterized by elevated expression of several valve-regulating transcription factors, including *Prox1, Nfat5,* and *Klf4* (**Table S4**), and it also more frequently expressed other key transcription factors (*Nfatc1*, *Gata2*, *Foxo1, Foxc2*, *Nr2f2* (COUPTF2)), and valve LEC markers (*Itga9, Cldn11, Piezo1)* (**Fig. 2c-e**, **Suppl. Fig. S4a**).

Overall, the molecular signatures of these cardiac LEC clusters were very similar to profiles previously reported for lymphatic subpopulations in other tissues, e.g. mouse skin^27^. Indeed, studies comparing LECs from different organs have found few organ-specific differences in transcriptomic signatures of healthy lymphatics^22^, despite accumulating evidence of organ-selective differences in developmental LEC origins^28^.

### Altered molecular profiles in cardiac ECs during pressure-overload-induced heart failure

Next, to define the transcriptome of cardiac ECs during pressure-overload, we repeated the sorting and sequencing procedures in 10 TAC-operated BALB/c mice at a stage of chronic heart failure^15^. After quality control, as described for healthy BALB/c samples, we obtained 1,455 EC transcriptomes. Following expression matrix normalization and batch-correction to integrate this dataset with healthy BALB/c EC transcriptomes, we found similar distribution post-TAC with cells clustering into three main populations: LECs (449 transcriptomes), BECs (838 transcriptomes), and vBECs (168 transcriptomes), as determined by Seurat graph–based clustering and visualization using UMAP (**Fig. 3a, b**). On average, in each cell, we detected 4,096 transcripts and 1,761 unique genes per cell (**Table S1**). The cell cycle phases of cardiac ECs post-TAC were similar as in healthy sham-operated hearts (**Suppl. Fig. S1b, c**). This finding of low EC proliferation is supported by our immunohistochemical data indicating rare Ki67^+^ cardiac ECs at 8 weeks post-TAC, as both angiogenic and lymphangiogenic responses triggered by the pressure-overload have abated in the chronic heart failure stage^15^.

**Fig. 3.**
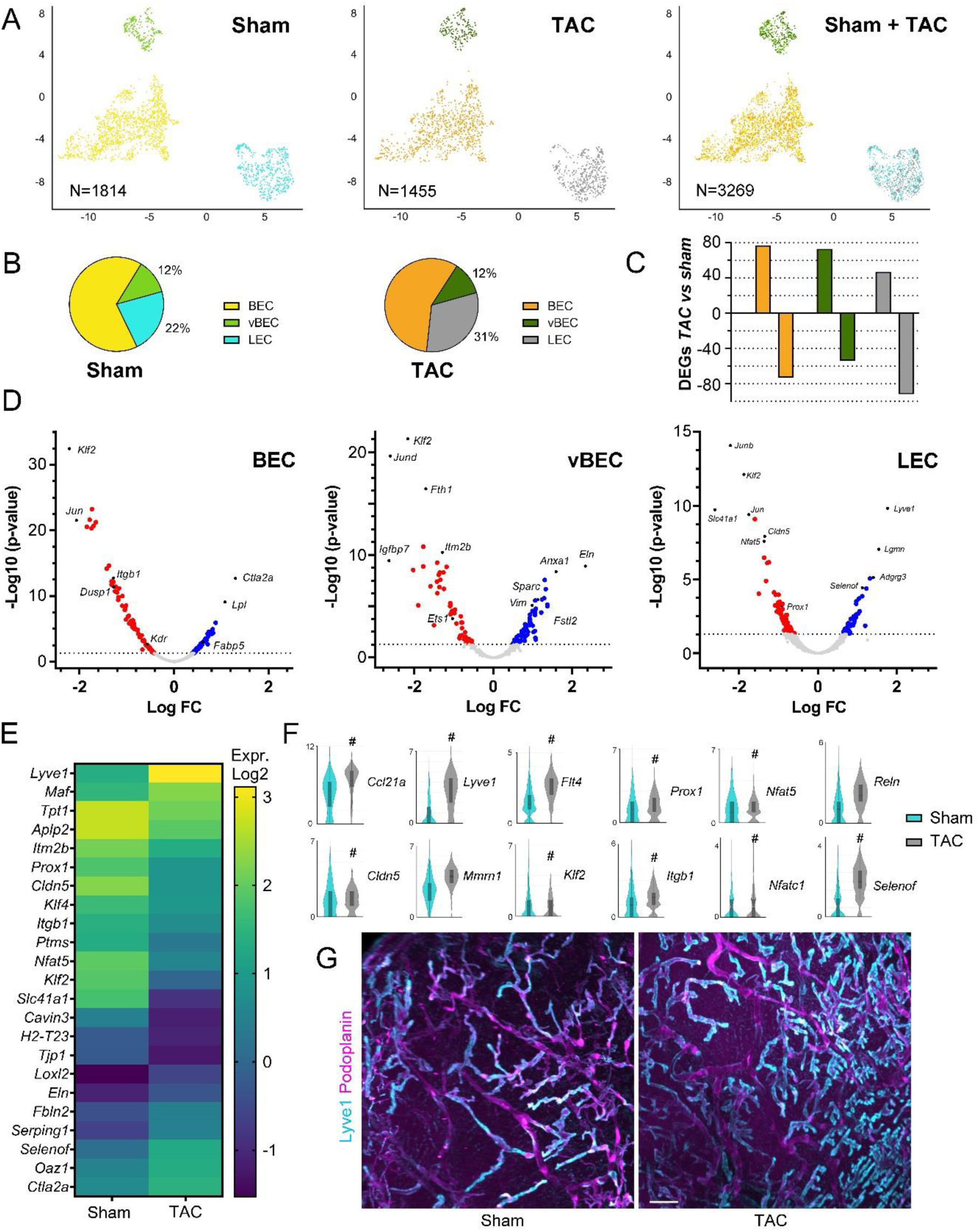
Molecular profile changes in the cardiac vasculature post-TAC in BALB/c. Cardiac ECs (*N=1,455*) from post-TAC BALB/c mice (*n=10*) clustered together with healthy cardiac ECs (*N=1814*) into three main EC populations: BEC, LEC, and vBEC (**a, b**). Numbers of DEGs (up- vs down-regulated post-TAC as compared to corresponding healthy clusters) in BEC, LEC and vBECs (**c**). Examples [volcano plot] of down- (*red*) and up-regulated (*blue*) genes in BEC, vBEC, and LEC clusters post-TAC, as compared to healthy hearts (**d**). For full list of DEGs identified post-TAC, see supplementary **tables S5, S6**, and **S7**. Examples of altered genes in cardiac LEC global cluster post-TAC [mean expression levels, *Log2 normalized read counts*: heatmap**, e**; violin plot, **f**]. Healthy LECs, *cyan*; post-TAC LECs, *grey*. Significant genes indicated (**#**) in panel **f**. Cardiac imaging (**g**) of healthy cardiac lymphatics versus expanded lymphatics at 8 weeks post-TAC. Lyve1, *cyan*; Pdpn, *magenta*. Scalebar 100 µm.

Next, comparing changes in gene expression induced by TAC in the different cardiac EC clusters, we identified 416 differentially-expressed genes (DEGs), including 139 in LECs (**Table S5**), 150 in BECs (**Table S6**), and 127 genes altered in vBECs (**Table S7**). Whereas the BEC cluster was perfectly balanced in the number of up- and downregulated genes, vBECs displayed slightly more upregulated genes (57% of DEGs), while more downregulated genes were found in LECs (66% of DEGs) (**Fig. 3c**).

In the LEC cluster post-TAC several immune-related genes were significantly altered. For example, *Lyve1*, *Ccl21*, and *Thy1* were upregulated, while *H2-T23* (MHC-I) was downregulated (**Fig. 3d-f**; **Table S5**). However, more LECs expressed *H2-D1* (94% of cells) and *H2-K1* (78% of cells) post-TAC, compared to healthy cardiac LECs (75% and 46% of cells, respectively) (**Suppl. Fig.S2b, Table S5**). More LECs also expressed *Serinc1* (Serine incorporator-1) post-TAC (78% vs. 48% of LECs), which encodes an enzyme stimulating synthesis of serine-derived lipids, such as S1P. However, the sphingosine biosynthetic pathway genes (kinases *Sphk1, Sphk2*, and lysase *Sgpl1*) were not altered, while cardiac LECs tended to increase *S1pr1* expression post-TAC (**Suppl. Fig. S3a**). In contrast, reduced *S1pr1* levels have been reported in inflamed dermal lymphatics in the setting of lymphedema^20^.

Further, we found that *Cldn5* was strikingly downregulated in LECs post-TAC (**Fig. 3d-f**), as was another tight junction component, *Tjp1* (**Suppl. Fig. S3a**), and several key lymphatic transcription factors, including *Prox1*, *Nfat5,* and *Nfatc1* (**Fig. 3d-f**, **Table S5**). However, the frequency expression of these lymphatic transcription factors, as well as *Nr2f2* and *Klf4* (Kruppel-Like factor-4), was increased, rather than decreased, in cardiac LECs post-TAC (**Suppl. Fig. S4b**).

Following pressure-overload, cardiac LECs also upregulated genes encoding extracellular matrix components, including *Eln* (Elastin), *Fbln2* (Fibulin-2), and *Loxl2* (Lysyl oxidase Like-2, a collagen cross-linking enzyme), as well as a proteinase inhibitor (*Serping1,* C1 inhibitor). Whereas the expression levels of *Reln* (Reelin) and *Aqp1* were not significantly increased, they were more frequently expressed (90% and 74% of cells expressed *Reln* and *Aqp1*, respectively, post-TAC, as compared with 62% and 45%, respectively, of healthy cardiac LECs).

Functional enrichment analyses, by Over Representation Analysis (ORA), was carried out to identify biological processes altered in cardiac LECs post-TAC. Top hits included (**Table S8**) (all with qvalue <3E^-19^): “*RNA splicing*” (GO:0008380, *padj* 3E^-35^), “*mRNA processing*” (GO:0050684, *padj* 3E^-19^), “*Actin reorganization*” (GO:0032956, *padj* 1E^-21^), and several processes related to cellular metabolism (e.g. GO:0015980, *padj* 7E^-19^). Further, “Ribosomal Biogenesis (GO:0042273, *padj* 1E^-3^) was upregulated post-TAC. Previous work has demonstrated that ribosomal biogenesis is increased in ageing BECs^29^, and the functional enrichment analyses indicated that stressed cardiac LECs during pressure-overload also may activate this process.

In line with the observed reduction of tight junction components in cardiac LECs post-TAC, KEGG pathway enrichment analysis identified several related down-regulated pathways, including “*Focal adhesion*” (mmu04510, *padj* 2E^-2^), and “*Fluid shear-stress*” (mmu05418, *padj* 2E^-2^).

Finally, reactome pathway enrichment analysis identified “*Cellular response to stress*” (R-MMU-2262752, *padj* 3E^-17^) and “*Cross-presentation of antigens*” (R-MMU-1236978, *padj* 7E^-19^) among key upregulated processes post-TAC.

In the BEC cluster, chronic pressure-overload led to upregulation of genes involved in lipid uptake and FA cytosolic trafficking (*Lpl*, *Fabp4, Fabp5*), and in regulation of extracellular matrix (*Timp4, Sparc)*. Further, while *Id3, S1pr1, Esam,* and *Icam2* expression levels were increased, *Ets1*, *Kdr, Tjp1,* and *H2-*D1 were reduced post-TAC (**Fig. 3d**, **Suppl. Fig. S2b, S3a**, **S5a**, For full list of DEGs, see **Table S6**). These profile changes are indicative of vascular EC activation, as previously described during pressure-overload-induced cardiac inflammation^30^. Investigating BEC subpopulations, we found similar subclustering of cardiac BECs post-TAC as in healthy hearts (**Suppl. Fig. S6a-d**). Of note, all BEC clusters displayed alteration of their gene expression profiles post-TAC (**Suppl. Fig. S3c, S6e**, For full list of DEGs, see **Table S9**). For example, the BEC1 cluster (enriched in capillary markers, including *Aqp1*) showed reduced expression levels post-TAC of *Ets1* and *Kdr,* and increased levels of *Timp4* and *Id3* (**Suppl. Fig. S6c, f**). *Lpl* was increased post-TAC in several distinct BEC subpopulations, while *Fabp4* was increased selectively in the *Notch1*-expressing BEC4 cluster (**Table S9**).

In the vBEC cluster, which represented around 15% of all blood vascular ECs in both healthy and failing hearts, we found many upregulated genes post-TAC (**Fig. 3c, d**, For full DEG list, see **Table S7**), including *Vim* (Vimentin; type III intermediate filament protein)*, Sparc, Aqp1, Selenof* (an endoplasmic reticulum-resident selenoprotein), and *Fstl1* (Follistatin-related protein-1). In contrast, the expression levels of *Nrp2*, *ApoE*, *H2-D1,* and *H2-K1* were reduced in vBECs post-TAC (**Suppl. Fig. S2b, S5b**). However, the frequency of expression of the two major MHC-I class molecules was increased post-TAC (96% and 89% of vBECs expressed, respectively, *H2-D1* and *H2-K1*), as compared to healthy vBECs (85% and 73% of cells expressing, respectively, *H2-D1* and *H2-K1*).

### Lymphatic subpopulation-selective molecular alterations and loss of lymphatic valves in failing hearts

Focusing on LEC subpopulations, we first established that the three LEC clusters identified post-TAC displayed similar molecular markers as in healthy mice, with no apparent unique cell population emerging post-TAC (**Fig. 4a**, **Table S4**).

**Fig. 4.**
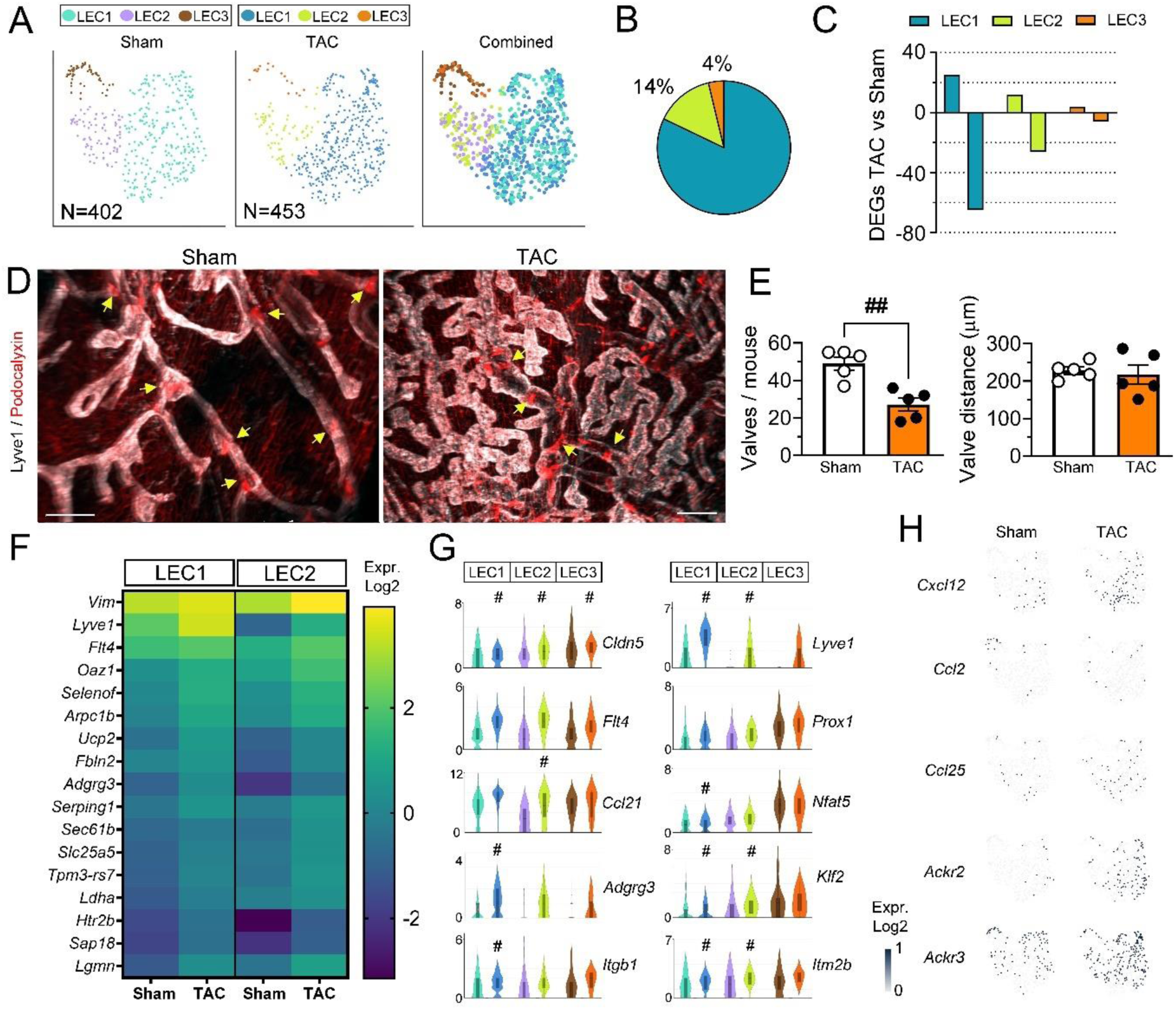
Modification of cardiac LEC subpopulations post-TAC in BALB/c. Cardiac LECs (*N=453*), obtained post-TAC in *BALB/c* mice (*n=10*), included three subpopulations: LEC1, LEC2, and LEC3. These cells clustered together with healthy cardiac LECs [umap, **a**]. The three LEC clusters post-TAC represented, respectively, 82%, 14%, and 4% of all LECs (**b**). Quantification of DEGs induced by TAC in the respective LEC clusters as compared to the corresponding healthy (sham) cardiac LEC clusters (**c**). Examples of cardiac lymphatic valves in healthy versus post-TAC mice (Lyve1, *grey*; Podocalyxin, *red*; lymphatic valves indicated by yellow arrows) (**d**). Scale bar, 200 µm. Quantification of lymphatic valves identified per mouse (n=5/group) in sham (white circles) and TAC (black circles) groups, and assessment of average lymphatic inter-valve distances in cardiac lymphatics (**e**). ##, p<0.01 Mann Whitney U-test. Examples of mean expression levels (*Log2 normalized read counts*) of altered genes [heatmap, **f**], [violin plot, **g**] in cardiac LEC subpopulations from healthy and post-TAC hearts. Significant genes indicated (**#**) for each cluster in **g**. For details see supplemental **table S10**. Comparison [umap, *normalized Log2 read counts*, **h**] of cardiac LEC expression of chemokines (*Cxcl12, Ccl2, Ccl25*), *Ackr2* and *Ackr3* in healthy and TAC mice.

We previously demonstrated that at 8 weeks post-TAC in BALB/c mice, the cardiac lymphatic network is massively expanded^15^, with selective overgrowth of Lyve1^high^ lymphatic capillaries (**Fig. 3g**). In agreement, our scRNAseq analyses of cardiac LEC subpopulations revealed an increase of the capillary LEC1 cluster, coupled to a reduction of precollector LEC2, and especially lymphatic valve LEC3 clusters (**Fig. 4a, b**). Indeed, whereas LEC1 and LEC2 represented 82% and 14%, respectively, the LEC3 cluster only included 4% of all LECs post-TAC. This represents a striking 4-fold decrease as compared to healthy sham-operated hearts (compare with **Fig. 2b**).

Our recent studies indicated that cardiac lymphatics appear dysfunctional post-TAC, as both cardiac immune cell infiltration and edema persisted despite lymphatic expansion^15^. Loss of lymphatic valves has been shown to lead to lymphatic transport dysfunction in different organs in mice^17^. However, no previous study has investigated whether cardiac lymphatic valves are altered in CVDs. As our scRNAseq data indicated striking rarefaction of the LEC3 population, we hypothesized that the number of lymphatic valves may be reduced in the heart post-TAC. To image these elusive structures, we performed wholemount imaging of Lyve1 together with Podocalyxin, which labels both lymphatic valves and blood capillaries (**Fig. 4d**). In healthy hearts, we found that lymphatic precollectors (low Lyve1-intensity, straighter vessel segments), but surprisingly also capillaries (defined as Lyve1^high^, tortuous segments), contained valves, with only the outermost tips of lymphatic capillaries completely devoid of valves. This organization is reminiscent of that described in dermal lymphatics^31^. In contrast, the expanded lymphatic capillary network established by 8 weeks post-TAC was almost completely devoid of valves (**Fig. 4d, e**). To further investigate lymphatic valve properties in the heart, we analyzed the distance between consecutive valves, finding an average intervalve distance of 200 µm (**Fig. 4e**). In the remaining valves in TAC-operated mice this valve distance was unaltered.

Next, we investigated whether lymphatic insufficiency post-TAC also may involve molecular profile changes in LEC capillaries (LEC1) impacting lymphatic cell-junctional organization or immune cross-talk. Analysis of LEC1, comparing TAC and sham, revealed 90 DEGs, including 65 downregulated genes (**Fig. 4c**, **Table S10**). Among these altered genes were *Lyve1*, which was increased, and the tight junction components *Cldn5* and *Tjp1*, which were decreased post-TAC (**Fig. 4f, g**). Another downregulated gene in the LEC1 cluster was *Aplp2* (Amyloid-like protein 2) (**Fig. 3e**, **Table S10**). The encoded protein stimulates endocytosis, notably of MHC-I molecules^32^, thus its reduction may influence MHC-I plasma membrane levels in lymphatics. Of note, this gene was also downregulated in BECs and vBECs post-TAC (**Tables S6, S7**).

Concerning potential mechanical stress-induced changes in the setting of chronic myocardial oedema post-TAC, we found that several mechanosensitive genes, including *Klf2* (Kruppel-Like 2), *Dtx1* (deltex E3 ubiquitin ligase-1), and *Itgb1* (Integrin β1), but not the ion channel *Piezo1*, were downregulated in cardiac LECs (**Fig. 4g**, **Tables S5**, **S10**). In lymphatics, *Klf2* has been shown to regulate pressure-induced LEC sprouting, in part through upregulation of *Dtx1*^33^, while *Itgb1* mediates shear-stress and interstitial fluid pressure-induced stimulation of lymphangiogenesis^34,35^.

Concerning lymphatic immune-attraction, we found that healthy cardiac LECs expressed low levels of *Cxcl12* and *Ackr3,* and these genes were not significantly altered post-TAC (**Fig. 4h**). *Ackr3* (*Cxcr7*) encodes an atypical chemokine receptor that binds and scavenges chemokines, e.g. Cxcl12. In contrast, other family members, *Ackr2* (*Ccr10*/D6) and *Ackr4* (*Ccr11*), which bind CC-type chemokines Ccl2 and Ccl21, respectively, were only rarely expressed (**Fig. 4h**). Of note, a recent scRNAseq study of inflamed dermal LECs reported a subpopulation of capillary lymphatics, denoted iLEC for immune-interacting cluster^27^. It was characterized by elevated expression of *Ptx3* (Pentraxin-3), and co-expressed *Aqp1, Stab1* (Clever-1*), Cd200, Mrc1* (CD206)*, Ackr2,* and *Plxnd1* (PlexinD1). In our dataset, we did not find such an “immune” capillary LEC subset emerging post-TAC. Indeed, *Ptx3* was only expressed by <6% of cardiac LEC1 cells in both healthy and post-TAC mice (**Suppl. Fig. S7a**). However, the frequency of *Ackr2* and *Mrc1* expression increased in LEC1 cluster post-TAC (*Ackr2*, 25% vs <5% of cells; *Mrc1*, 41% vs. 12% of cells in healthy hearts). Nevertheless, cells expressing these proposed iLEC markers did not display convincing clusterisation within the LEC1 population (**Suppl. Fig. S7a, b**).

In the LEC2 cluster, the 38 DEGs identified post-TAC (**Table S10**) included 12 upregulated genes, notably *Lyve1* and *Ccl21*, and 26 downregulated genes, including *Cldn5, Klf2*, *Cavin3* (Caveolae associated protein-3), *Itm2b (*Amyloid-binding Integral Membrane Protein-2b), and *Tgfrb3* (Transforming growth factor receptor β3).

DEG analyses in the LEC3 cluster lacked sensitivity due to the limited number of cells present post-TAC. Among the few identified upregulated genes was *Cavin2* (**Table S10**). In contrast, there was no significant change in expression levels or frequency of expression of valve-inducing transcription factors (*Prox1, Nfat5, Foxc2, Nfatc1, Gata2*) in the remaining LEC3 cluster post-TAC (**Fig. 4f, g**, **Suppl. Fig. S4a**).

### Few modifications of cardiac ECs during pressure-overload in C57 strain

To determine whether loss of lymphatic valves and dysregulation of cardiac EC expression profiles also occurred in a setting of less severe pressure-overload-induced cardiomyopathy, we repeated the cardiac EC sorting and scRNAseq in C57BL6/J (C57) adult female mice. We previously showed that in this strain of mice chronic pressure-overload, by the same surgical procedure used in BALB/c, leads to pathological LV hypertrophy, accompanied by milder cardiac inflammation and less LV dysfunction at 8 weeks post-TAC^15^. We also reported in these mice limited expansion post-TAC of cardiac lymphatics, which seemed to retain more physiological-like organization, as evaluated by wholemount imaging. Sorting cardiac single cell suspension from 10 mice per group, we obtained, after quality control filtering (removing genes expressed by <10 cells; and removing cells expressing <300 genes, or >10% mitochondrial-encoded genes), 634 ECs from sham-operated mice and 892 ECs from TAC-operated C57 mice. On average, in each cell, we detected 3500-5400 transcripts and 1600-2200 unique genes per cell (**Table S1**). The transcriptomes clustered into LECs, BECs, and vBECs, similar as described for BALB/c, with 142 genes distinguishing these cardiac EC populations (**Fig. 5a, b, e**).

**Fig 5.**
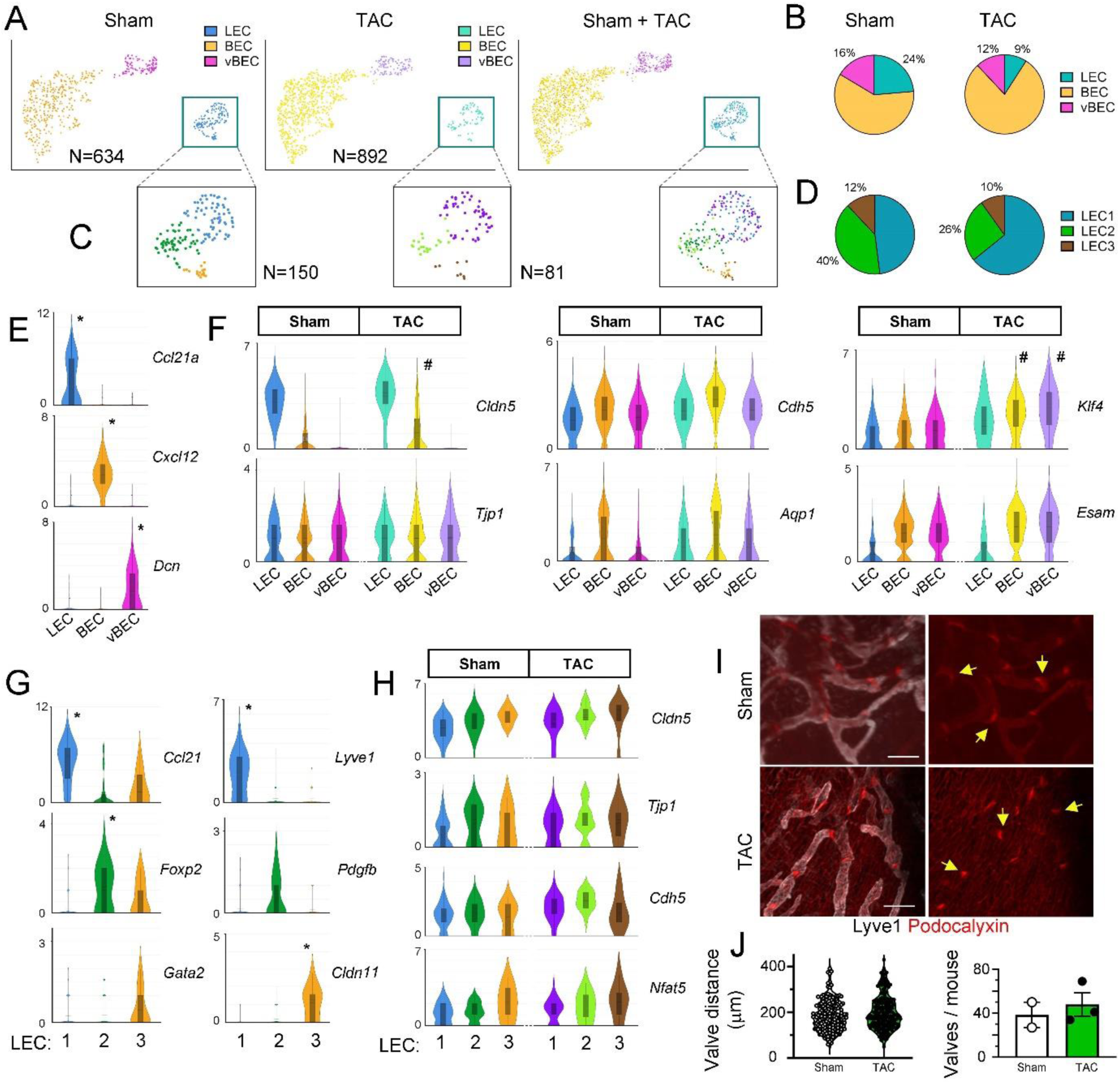
Limited impact of chronic pressure-overload on cardiac ECs in C57 mice. Unsupervised clustering of cardiac ECs, from healthy or post-TAC C57 hearts (*n=10* mice per group, N=634 and N=892 transcriptomes, respectively) revealed three main EC populations (**a** [umap], **b**): BEC, LEC, and vBEC. Cardiac LECs (*N=150* healthy*; N=81* post-TAC) clustered into three main lymphatic subpopulations: LEC1, LEC2, and LEC3 (**c** [umap], **d**). Examples [violin plot, *Log2 normalized read count*] of cardiac EC marker genes (**e**), and barrier-relevant genes and DEGs in TAC vs sham C57 mice (**f**). Significantly-enriched marker genes are indicated (**\***), and genes significantly upregulated post-TAC are indicated (**#**) for respective clusters. For full list of DEGs, see supplemental **table S11**. Examples [violin plot, *Log2 normalized read count*] of LEC cluster markers (**g**), and expression of barrier-relevant genes in sham and TAC C57 hearts (**h**). Cardiac wholemount (**i**) visualization of lymphatic valves in healthy (*top*) versus post-TAC (*bottom*) C57 mice (Lyve1, *grey*; Podocalyxin, *red*; lymphatic valves indicated by yellow arrows). Scale bar, 100 µm. Quantification of lymphatic valves per mouse in sham (*white circles*) and TAC (*black circles*) groups, and assessment of intervalve distances in cardiac lymphatics (**j**).

Next, differential gene expression analyses, within clusters and between groups, revealed that chronic pressure-overload induced only very minor shifts in cardiac EC molecular profiles, with totally 34 DEGs identified in cardiac ECs post-TAC (**Table S11**). This included upregulation of *Cldn5* in BECs, and *Klf4* in both BEC and vBEC clusters (**Fig. 5f**).

Within the global cardiac LEC population in healthy C57 mice, we identified three small, but distinct, LEC clusters (**Fig. 5c, d**). Of note, significantly fewer cardiac LECs were recovered from C57 mice, either sham (N=150 single-cell transcriptomes) or TAC (N=81 single-cell transcriptomes), as compared to BALB/c. Nevertheless, identified markers of cardiac LECs in healthy C57 mice, and of its subpopulations, LEC1 (48% of all LECs), LEC2 (40% of all LECs), and LEC3 (12% of all LECs), mirrored those found in BALB/c (**Fig. 5e, g**). For example, the C57 LEC3 cluster expressed lymphatic valve marker genes (e.g. *Cldn11*, *Nfat5, Gata2*). We only detected 3 DEGs in cardiac LECs post-TAC, and none of them influenced lymphatic chemokines, immune adhesion molecules, or barrier components (**Fig. 5h**, **Table S11**). Indeed, the LEC cluster post-TAC in C57 mice had very similar molecular profile as healthy LECs, in strong support of a more physiological-like state of cardiac lymphatics during pressure-overload in C57, as compared to in failing BALB/c hearts.

Enrichment analysis (ORA) of LEC transcriptomes revealed upregulation post-TAC of processes including “*positive regulation of cell activation*” (GO:0050867, *padj 9E^-3^*), “*positive regulation of leukocyte activation*” (GO:0002696, *padj 3E^-2^*), and “*positive regulation of B cell proliferation*” (GO:0030890, *padj 3E^-2^*).

Concerning lymphatic valves, we found that the frequency of valvular LECs in healthy mice were comparable between C57 and BALB/c strains, and strikingly we only found a minor reduction in the frequency of LEC3 cells post-TAC (**Fig. 5d**). Our wholemount analyses further confirmed that lymphatic valves were essentially retained after pressure-overload in C57, with additionally no change in cardiac lymphatic intervalve distances (**Fig. 5i, j**).

### Validation of altered molecular profiles post-TAC

Next, we set out to verify, at the protein level, some of the identified cardiac lymphatic DEGs in BALB/c during pressure-overload. We focused our analyses on lymphatic: 1) Lyve1 and Ccl21 expression levels; 2) cell junctional components (Cldn5); and 3) Reelin levels.

Wholemount confocal imaging and flow cytometry both confirmed a striking increase in cardiac lymphatic Lyve1 levels post-TAC (**Fig. 6a, b**). Similarly, *Ccl21* upregulation post-TAC was confirmed by quantitative analyses of peri-lymphatic Ccl21 deposition in cardiac tissue sections, as recently reported^15,16^. Thus, active Ccl21-mediated lymphatic recruitment of CCR7^+^ immune cells, and/or Lyve1-mediated transcytosis of hyaluronic acid-decorated immune cells^9^, is likely to be increased post-TAC in BALB/c.

**Fig 6.**
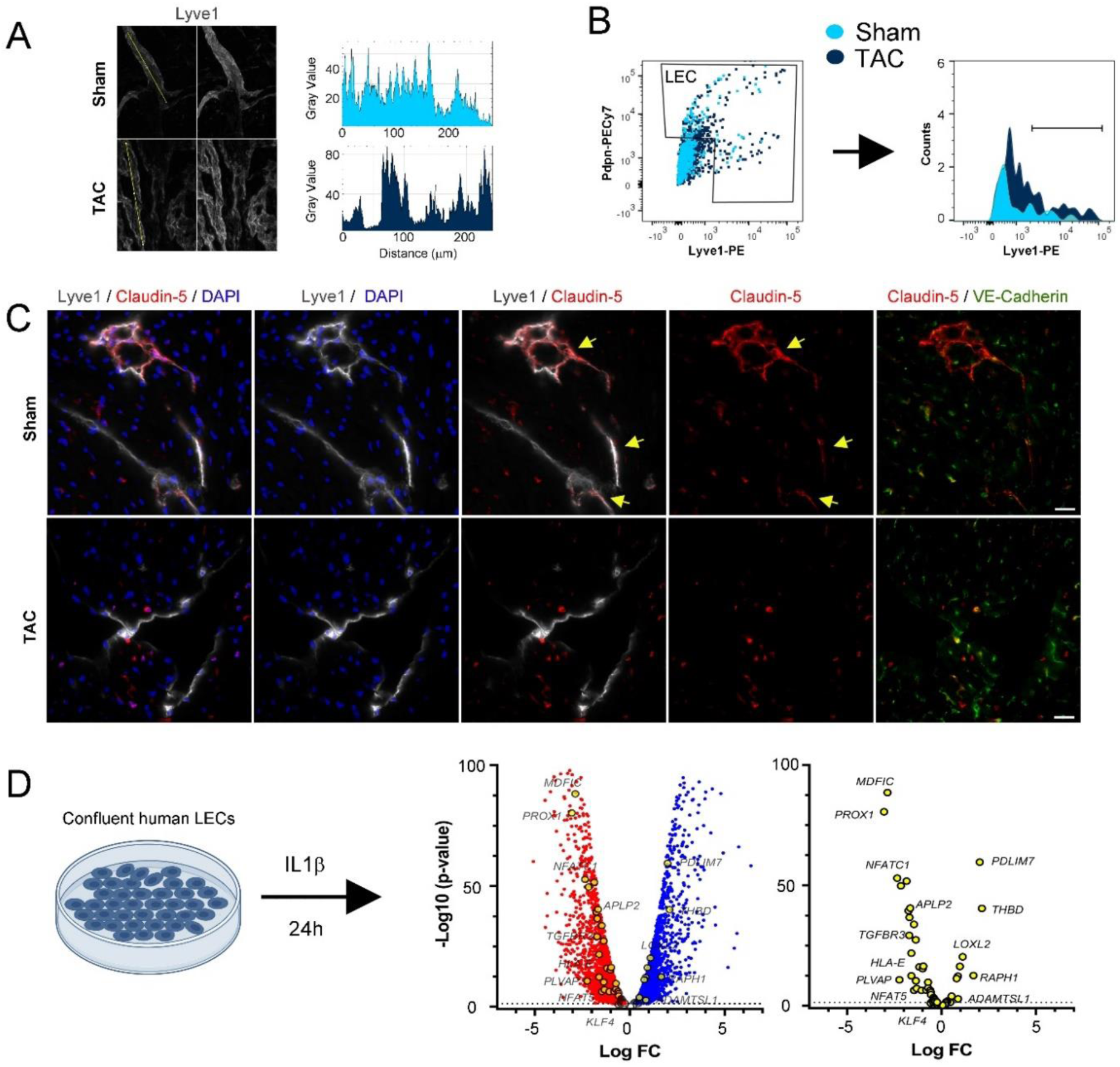
Lymphatic profile changes induced by cardiac inflammation. Confocal imaging (**a**) used for quantification of Lyve1 intensity in cardiac wholemounts in healthy and post-TAC BALB/c. Yellow line outlines the lymphatic segment analyzed for signal intensity using Image J. Flow cytometric assessment (**b**) of Lyve1 expression in isolated cardiac LECs from healthy and TAC BALB/c. Examples (**c**) of Claudin-5 (*red*) expression in healthy and post-TAC cardiac lymphatics (Lyve1, *grey*) and blood vessels (VE-Cadherin, *green*). Scalebar 20 µm. Arrows point to Claudin-5-expressing lymphatics. Examples [volcano plot] of DEGs (**d**), either down- (*red)* or up-regulated (*blue)* genes, following IL-1β stimulation of confluent human LECs *in vitro.* Genes similarly altered by IL-1β in human LECs, as observed in BALB/c cardiac LECs post-TAC, are high-lighted (yellow circles). For details see supplementary **table S12** and **Fig. S8**.

In agreement with potent down-regulation of *Cldn5* observed in LECs during chronic pressure-overload, our immunohistochemical analyses revealed striking reduction of lymphatic Claudin-5 levels in BALB/c hearts at 8 weeks post-TAC (**Fig. 6c**). In contrast, while our scRNAseq data suggested slight upregulation of *Reln*, we could not detect Reelin by immunohistochemistry in either healthy or post-TAC mouse hearts.

Next, we speculated that part of the dysregulated molecular profile observed in lymphatics post-TAC in BALB/c could be linked to elevated cardiac proinflammatory cytokine levels in failing hearts, notably IL1β^15^. To address this possibility, we stimulated human LEC primary cultures *in vitro* with IL1β for 24h followed by bulk RNAseq to establish inflammation-induced DEGs (**Table S12**). Among the 135 DEGs identified in the LEC cluster post-TAC (**Table S5**), we found 50 genes that displayed similar changes in IL1β-treated human LECs (**Fig. 6d, Suppl. Fig. 8**). This suggests that a considerable part of the transcriptomic changes observed in lymphatics during pressure-overload may be induced by cardiac proinflammatory cytokines, including IL1β. Specific examples of shared DEGs include upregulation of *Pdlim7* (PDZ And LIM Domain 7), *Selenof*, *Loxl2*; and downregulation of *Aplp2*, and of several lymphatic transcription factors (*Prox1, Nfat5*, *Nfatc1)*, and of *Plvap* (Plasmalemma vesicle–associated protein). Of note, while *Tjp1* was reduced by IL1β in human LECs, *Cldn5* expression was not altered.

In human LECs we also found that IL1β upregulated *CD274* (PDL1) and several MHC genes. This suggests that LEC-mediated antigen-presentation during cardiac inflammation may drive immune tolerance rather than T cell activation, similar as reported in tumor models^21^. However, *Cd274* expression was not significantly increase in cardiac LECs post-TAC. Indeed, both our scRNAseq and immunohistochemical analyses showed low expression levels of Pdl1 post-TAC (**Suppl. Fig. S7c, d**).

## Discussion

### Distinguishing traits of cardiac lymphatic subpopulations in health and disease

To the best of our knowledge, this study represents the first targeted molecular analysis of cardiac LEC subpopulations. As lymphatics represent a rare cell population in the heart, being restricted in rodents to only the outer cardiac layers, an approach of LEC enrichment through FACS sorting was required. Of note, we only obtained around 200-300 cardiac LECs per mouse in BALB/c, and even fewer cells in C57 mice, by FACS, and our scRNAseq thus included cells derived from 10-20 mice per group to yield sufficient numbers to allow lymphatic subpopulation analyses. Previously published expression analyses in cardiac LECs include: 1) targeted qPCR in C57 mice^5^; 2) bulk RNAseq in C57 mice^13^; and 3) scRNAseq of sorted PECAM1^+^ vascular ECs in human developing hearts^36^, and in adult C57 mice^22^, including one small LEC cluster each. Importantly, our study of LECs from healthy BALB/c and C57 hearts showed overall similar molecular profiles as these previous studies. However, our work goes well beyond these datasets by: 1) providing insight into what differentiates lymphatic capillaries from precollector and valve LECs in the heart; and 2) allowing cellular dissection of which compartment(s) of the lymphatic tree is affected during myocardial inflammation triggered by pressure-overload in a strain-dependent manner. Our molecular studies revealed that, different from our wholemount imaging of cardiac lymphatics (which suggested lower Podoplanin levels in capillaries as compared to precollectors), the gene expression levels of *Pdpn* were comparable between LEC subpopulations. In contrast, both gene expression and protein levels of Lyve1 were higher in cardiac lymphatic capillaries (LEC1) as compared to precollectors (LEC2), in line with findings from other tissues^27^. Further, different from our expectations, our study did not allow identification of new molecular markers specific for precollector lymphatics. Indeed, although the cardiac LEC2 cluster more frequently expressed *Pdgfb* and *Foxp2*, their expression levels were not homogenous, and would not allow unequivocal identification of precollectors in histological sections. Moreover, although several valve LEC-specific markers were readily identifiable, corresponding to markers described previously for lymphatic valves in other tissues^27^, the scarcity of valve LECs, notably in CVD settings, precludes analyses of these cells in tissue sections. Thus, it is still necessary to perform 3D imaging to differentiate, by immunohistochemistry, between the different segments composing the lymphatic vasculature of the heart.

### Molecular impact of inflammation on cardiac lymphatic functions

Inflammation has been reported to induce lymphatic dysfunction in many different organs, as pro-inflammatory mediators, including IL1β, act on lymphatic muscle cells (LMCs) to reduce lymph drainage^37^. Conversely, in the heart, where lymphatic drainage essentially depends on the force of cardiac contractions^38^, the negative inotropic effects of IL1β^39^ may contribute, indirectly, to insufficient cardiac lymphatic transport in CVDs. In addition, our scRNAseq analyses, coupled to our *in vitro* study in human LECs, revealed that the pro-inflammatory micro-environment in CVDs also may directly impact lymphatic function by altering LEC expression of genes regulating: 1) cell junctional organization/ stability impacting uptake and/or transport of fluids and macromolecules (*Cldn5, Tjp1, Plvap*), 2) immune cell attraction and adhesion (*Ccl21a*, *Lyve1, Thy1, Ackr3*); and 3) immune-modulation (*H2-T23, Fgl2, Aplp2, Cd274*).

Previous studies have identified an additional important role of cardiac lymphatics, related to production of Reelin^11^. Reelin is a large secreted glycoprotein that binds to several cell receptors, such as ApoE-R2, VLDLR, integrins, and ephrins. It is released by lymphatics under conditions of ischemia and/or inflammation, and has been shown in mice to act as a lymphangiocrine factor stimulating cardiomyocyte survival post-MI^2,11^. However, in our study, the non-significant upregulation of Reelin in cardiac LECs post-TAC could not be confirmed on tissue sections using immunohistochemistry.

#### Reduced lymphatic junctional stability during pressure-overload?

We observed reduction post-TAC of *Cldn5* and *Tjp1* in cardiac LECs only in BALB/c mice. The reduction of Claudin-5 was confirmed by immunohistochemistry in cardiac sections. Reduced expression of tight junction components may influence lymphatic junctional organisation (button/zipper junction). In lymphatic capillaries, looser cell-cell adhesion may facilitate fluid uptake during myocardial oedema, while loss of junctional stability in precollectors would lead to fluid leakage and hence reduced transport capacity. However, a recent study reported that LEC *Cldn5* deletion did not cause complete breakdown, but only remodeling, of lymphatic junctions^40^. In agreement, we previously reported no striking differences in the frequency of lymphatic capillary button-junctions post-TAC^15^, while they were clearly reduced, and replaced by zipper-junctions, post-MI^12^. Thus, the functional impact of low Cldn5 levels in cardiac LECs post-TAC remains to be determined. Of note, while IL1β altered the expression of *TJP1* and several claudins in human LECs, *CLDN5* was unaltered. This indicates that processes other than inflammation may underly loss of Cldn5 in cardiac lymphatics during heart failure. In addition, we found down-regulation of *Plvap* in cardiac lymphatics (global LEC and LEC1 clusters) post-TAC and in IL1β-treated human LECs. This gene is a marker of fenestrated ECs, and has been proposed to restrict immune cell passage through lymphatics^41^, hence down-regulation of *Plvap* post-TAC may result in increased cardiac lymphatic permeability.

Our scRNAseq data can be compared with cardiac lymphatic transcriptomes, obtained by bulk RNAseq, in another model of chronic pressure-overload induced by Angiotensin-2 (AngII)^13^. This model, performed in male C57 mice, resulted in chronic hypertension, moderate cardiac hypertrophy and dysfunction, and rarefaction of cardiac lymphatics, which all were improved by VEGF-C_C156S_ therapy. In contrast, we and others have previously reported maintenance, but not rarefaction, of cardiac lymphatics post-TAC in C57 mice, with accelerated disease after anti-VEGFR3 treatment^5,15^. Different from our findings in BALB/c post-TAC, but in line with our results in C57, Song *et al.*^13^ did not report reduced expression of genes relevant for lymphatic barrier, including *Cldn5* and *Tjp1*. Moreover, *Esam, S1pr1,* and *Jam2* levels appeared increased in their model of AngII-induced pressure-overload^13^ (**Suppl. Fig. S9**).

#### Enhanced cardiac lymphatic immune cell attraction and adhesion in pressure-overload?

Our study revealed striking upregulation, at both gene and protein levels, of the main lymphatic chemokine, Ccl21, post-TAC only in BALB/c. In addition, *Lyve1* and *Thy1* were upregulated. In contrast, previous studies of cardiac LEC in C57 have demonstrated that *Ccl21* gene expression was either not altered or reduced following pressure-overload^5,13^; while Lyve1 levels were reduced post-TAC^5^, but unaltered in the AngII model^13^.

We were surprised by the limited number of chemokines expressed by cardiac LECs in our study. Indeed, no other chemokine, besides *Ccl21a*, was differentially-expressed in cardiac lymphatics post-TAC. This may reflect poor sensitivity of scRNAseq approaches for weakly-expressed genes. In contrast, bulk RNAseq of cardiac LECs^13^ or lymph node LECs^42^, have revealed a significantly wider range of lymphatic chemokines in both health and disease. In agreement, using bulk RNAseq, we found potent upregulation of many chemokines (e.g. *CCXL1, CXCL8, CCL2*, *CXCL6*) in human LECs following IL1β stimulation. Previous studies have reported upregulation of *Cxcl12* in cardiac LECs in AngII-induced, but not TAC-induced, pressure-overload in C57 mice^5,13^. In parallel, recent scRNAseq of cardiac-infiltrating immune cells post-MI and post-TAC in C57 mice have demonstrated that while Cxcr4 (the receptor for Cxcl12), mainly is expressed by myeloid cells, including *Trem2*-expressing macrophages^43^, *Ccr7* (the receptor for Ccl21), is preferentially expressed by B and T cells^44^. We conclude that in our pressure-overload model in BALB/c, cardiac lymphatics are well-equipped to drain cardiac-infiltrating immune cells, including Ccr7-expressing B and T cells and hyaluronan-coated CD44-expressing myeloid cells interacting with Lyve1. In agreement, we previously reported low cardiac B cell levels post-TAC in BALB/c, while intriguingly CD4^+^ T cell levels remained significantly elevated as compared to healthy mice^15^.

#### Immune-modulation by cardiac LECs during cardiac inflammation?

Cardiac LEC activation by the pro-inflammatory micro-environment established during chronic pressure-overload, including elevated IL1β, could potentially influence lymphatic immune-modulation beyond immune cell attraction and adhesion. As mentioned above, *Ptx3* has been proposed as a marker of an iLEC lymphatic subpopulation^27^. Moreover, *Ptx3* was expressed in cardiac LECs from human fetal hearts^36^. In contrast, we did not find a cardiac LEC subpopulation homogenously expressing *Ptx3*. Indeed, this gene was expressed at low levels and frequency in cardiac LECs. Further, in our study LEC1 expression levels of additional markers proposed for iLECs (*Mrc1, Plxnd1, Aqp1, Ackr2, Cd200)* were also not altered post-TAC, although the frequency of expression of these markers did increase. Nevertheless, we could not identify an iLEC subpopulation within the LEC1 cluster. We speculate that the difference noted between our *in vivo* study and that of *Petkova et al.*^27^ in part may reflect differences in LEC proliferative status, with active expansion of dermal LECs occurring in their model, as compared to the more quiescent state of cardiac lymphatics at 8 weeks post-TAC in our study. It is also possible that differences in levels of pro-inflammatory immune-mediators could be involved. Indeed, our *in vitro* study demonstrated that acute IL1β stimulation increased the expression of several iLEC markers (*PTX3*, *PLXND1, CD274*, *ACKR3*, *CD200)* in human LECs.

We also found in cardiac LECs post-TAC in BALB/c mice altered expression of genes relevant for antigen-presentation, including more frequent expression of several MHC-I molecules and reduced *Aplp2* expression levels. Further studies are needed to assess the potential immune-modulatory / tolerogenic role of cardiac LECs in different CVD settings.

## Conclusion & Perspectives

Our study, together with previous reports^5,13^, indicates that cardiac lymphatics are more activated, on a cellular level, following pressure-overload in BALB/c, as compared to in C57 mice. This is in line with more potent cardiac lymphangiogenesis in the former^15^. Indeed, it appears that the less expanded lymphatic network in hypertrophic C57 hearts, irrespective of the pressure-overload model (AngII^13^ or TAC^5^), remains more physiological-like, as compared to the structurally- and molecularly-altered expanded lymphatics found in failing hearts in BALB/c. Of note, cardiac levels of several pro-inflammatory cytokines are higher post-TAC in BALB/c, compared to in C57^15^. This difference may contribute not only to more potent lymphatic expansion in the former, given the emerging pro-lymphangiogenic impact of cardiac IL1β^16^, but also to dysregulation of LEC molecular profiles during chronic pressure-overload-induced heart failure (**Fig. 7**). Indeed, our *in vitro* data in human LECs provided compelling evidence for a key role of IL1β in mediating cardiac inflammation-induced alterations of lymphatic transcriptomes. In agreement, we recently reported that IL1β blockage prevented pressure-overload-induced lymphatic Ccl21 upregulation^16^. However, it must be pointed out that the treatment also reduced cardiac lymphangiogenesis and did not prevent heart failure development in this model^16^, indicating that other changes in the cardiac micro-environment are involved in mediating lymphatic molecular remodeling in heart failure.

**Fig 7.**
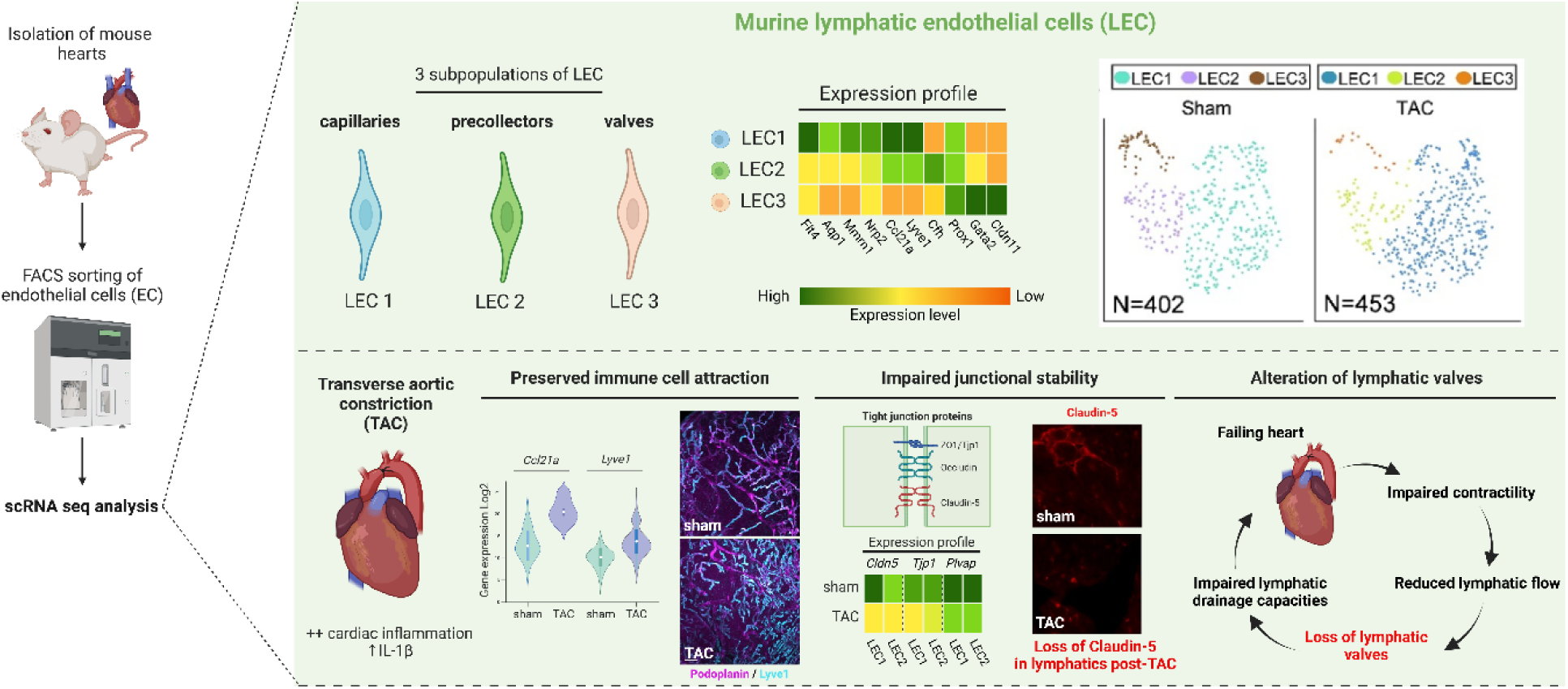
Overview of changes to cardiac lymphatics in Heart Failure. Our study uncovered distinct molecular profiles of cardiac lymphatic subpopulations, including capillary, precollector, and valve LECs. Chronic pressure-overload, induced by TAC surgery, led to structural, cellular, and molecular changes in cardiac lymphatics. This included upregulation of genes involved in immune cell attraction, downregulation of genes involved in junctional stability, and rarefaction of valvular LECs. We speculate that cardiac contractile dysfunction during development of heart failure leads to reduced lymphatic transport, causing loss of shear-stress signals necessary for maintenance of lymphatic valves. Rarefaction of lymphatic valves, together with inflammation-induced alterations of LEC molecular profiles, results in further lymphatic dysfunction contributing to poor resolution of edema and inflammation, promoting progression of heart failure (created using BioRender).

Concerning new molecular targets to enhance lymphatic function in CVDs, our current study revealed as the most notable change a radical loss of valves in cardiac lymphatics post-TAC. Single-cell analyses of the few remaining cells in the LEC3 cluster post-TAC in BALB/c lacked sensitivity to reveal potential dysregulated pathways, beyond loss of *Cldn5,* to explain this selective rarefaction. Of note, valve formation in lymphatics is strongly influenced by mechanosensitive (*Piezo1, Klf2, Klf4)* and/or calcineurin pathways (*Nfat)*^28,45^. Intriguingly, we found that several relevant genes (*Klf2*, *Dtx1*, *Nfat5*, *Nfatc1, Itgb1*) were reduced in BALB/c cardiac LECs post-TAC. Based on these alterations, we speculate that reduced lymphatic flow in failing hearts may be involved in lymphatic valve dropout, which in turn will reduce lymphatic drainage capacity. However, given that the reduction in cardiac LECs post-TAC of the *Klf2* gene was a feature shared with cardiac BECs and vBECs, it is also possible that loss of mechanosensitive pathways may reflect a global adaptation of cardiac ECs to chronically-increased myocardial fluid pressure and vascular shear-stress during TAC-mediated pressure-overload.

In conclusion, our study demonstrated severe changes in cardiac lymphatics during pressure-overload-induced heart failure, with in contrast very few alterations noted during pressure-overload-induced pathological hypertrophy. Future studies, perhaps at earlier timepoints before loss of valves, are required to provide tractable targets to restore cardiac lymphatic function in heart failure.

## Material and Methods

### Study approval

Animal experiments were approved by the regional Normandy University ethical review board Cenomexa according to French and EU legislation (APAFIS #23175-2019112214599474 v6; APAFIS #32433-2022070712508369 v2). A total of 60 BALB/c and 30 C57BL6/J female mice, surviving TAC or sham-operation were included in this study.

### Mouse *in vivo* model

Cardiac lymphatics were investigated in sham-operated or TAC-operated adult (8-week old) female BALB/c and C57BL6/J mice (Janvier Laboratories, France). Briefly, a minimally invasive method was used to constrict the aortic arch, using double-banding across a 26G needle as described^15^. Mice were anaesthetized by intraperitoneal injection of ketamine (100 mg/kg Imalgene^®^) and xylazine (10 mg/kg Rompun^®^ 2%, Bayer Health Care). Buprenorphine (50 µg/kg, Buprecare^®^, Axcience) was injected subcutaneously 6 hours after surgery and twice per day until 3 days post-operation. Euthanasia was performed by pentobarbital overdose (100 mg/kg Euthoxin) at 8 weeks post-TAC.

### Cardiac single cell sorting, sequencing, and analyses

Cardiac single cell suspensions were sorted by FACS (ARIA II, BD Biosciences). BECs were defined as live CD45^-^/CD31^+^/Lyve1^-^ cells, whereas LECs were identified as CD45^-^/CD31^+^/Lyve1^+^/Pdpn^+^ cells. A 1:3 mix of cardiac LECs and BECs were included in each sample prepared using the 10X Genomics Chromium Next Gem Single Cell 3’ kit. For each group 1-2 reactions were performed using each time cells pooled from 10 mouse hearts. Samples were sequenced using Illumina NextSeq550 at an estimated sequencing depth of 20 000 reads per cell. Raw FASTQ files were generated and demultiplexed from BCL data using bcl2fastq (v2.20.0). Quality control with FastQC (v0.11.9) indicated >98% perfect index and 86% of bases >= Q30, requiring no treatment, with the exception of BALB/c sham1 sample trimmed using FASTP (v0.20.0) to remove adapters, poly-A, and poly-G present in Reads. Reads were aligned to the mouse reference genome (GRCm39.111) using STARsolo^46^ RNAseq aligner (v2.7) to produce raw UMI count matrix. Data filtering, normalization, dataset integration, principal component-based unsupervised clustering, and differential analysis were performed using the Seurat package (version 5.0.2)^47^ in R (version 4.1.2). Functional enrichment analysis was performed with the R package clusterProfiler (v4.10.2). In BALB/c mice, among 3908 total sequenced cells, 3436 cardiac cells remained post-filtering, including 1932 cells from sham-operated BALB/c controls and 1504 from post-TAC mice. In C57BL6/J mice, among 4178 total cells, 1577 cardiac cells remained post-filtering, including 665 and 912 cells from sham and post-TAC mice, respectively. Processed data was exported for visualisation using CellLoupe (Loupe Browser 8.1.2, *10x Genomics*) to generate UMAP and violin plots. Heatmaps and Volcano plots were generated based on Seurat-identified mean gene expression levels (plotted as log2 transformed normalized read counts), and log2 fold-change (FC) and adjusted p-values, respectively. Statistical analyses were performed to determine differences in mean gene expression levels: 1) between clusters of cardiac ECs in either healthy or post-TAC mice, to identify marker genes; 2) between TAC and sham groups, within specific clusters, to identify impact of pathology. Only genes with a mean normalized read count >0.3 were included in differential analyses. Results of DEG analysis and functional enrichment analysis are presented in supplementary tables S2-S11. For details see *Suppl. Methods*.

### Validation by immunohistochemistry

Cardiac sections were analyzed by immunohistochemistry to determine lymphatic expression of target genes identified by scRNAseq. Whole mount-staining of cardiac lymphatics was performed, using a modified iDISCO^+^ clearing protocol, for imaging by light sheet and confocal laser scanning microscopy, as described^15^. For details see *Suppl. Methods*.

### Validation by *in vitro* culture of human LECs

Briefly, human LECs (Clonetics™ HMVEC-dLy, *CC-2812*) were grown to confluence according to recommendations of the supplier. After overnight serum-starvation (1% FCS), cells were exposed to 20 ng/mL recombinant human IL-1β (PeproTech) for 24h. RNA was extracted (Single Cell RNA Purification Kit, Norgen) for mRNA IlluminaSeq by Biomarker Technologies (BMK, Germany). For details see *Suppl. Methods*.

### Statistics for bulk RNAseq and immunohistochemistry

Bulk RNAseq data from human LEC cultures was analyzed for gene expression levels and DEGs by BMK using DESeq2 bioconductor package (Wald test). Data is reported as Fpkm normalized reads and as fold-change (FC) of control unstimulated cells. Statistical analyses for immunohistochemical data were performed using GraphPad Prism software. Data are presented as mean ± s.e.m. Comparisons of two independent groups were performed using either Student’s two-tailed t-test for groups with normal distribution, or alternatively by Mann Whitney U test for samples where normality could not be ascertained based on D’Agostino & Pearson omnibus normality test. For details see *Suppl. Methods*.

## Acknowledgement

We thank Dr Maria Goes (Max-Planck-Institute for Heart and Lung Research) for helpful suggestions for Ccl21 wholemount staining; Ms Marine Panza (Univ Rouen) for assistance with immunohistochemistry; and Mr Clément Lecler (Univ Rouen) for assistance with library generation for scRNAseq.

## Contribution

C.H and T.L performed single-cell preparations for scRNAseq with guidance by G.R and S.C; mRNA sequencing was performed by C.D; scRNAseq data analysis pipeline was designed and executed (including generation of expression count matrices and Seurat analysis) by B.B and C.T, with guidance by C.B and H.D; immunohistochemistry, image acquisition, and analyses were performed by A.D, O.L, C.H, C.V, D.G, and D.S; cell culture experiments, including RNA extraction for bulk RNAseq was performed by O.L; surgical mouse model was carried out by M.V; A.Z, H.D, and V.T provided critical feedback on experimental design and on the manuscript; E.B, C.H., and T.L designed the study, analyzed results, and prepared the manuscript draft. All authors approved of the final version of the manuscript.

## Data Availability

The original data included in this study has been deposited in GEO (GSE289738, *Effect of interleukin-1beta on confluent human lymphatic endothelial cells;* GSE290576 *Effect of pressure-overload on cardiac endothelial cells*). Data (**Suppl. Fig. 9**) reproduced from Song, L. et al. *Lymphangiogenic therapy prevents cardiac dysfunction by ameliorating inflammation and hypertension*. Elife 9, e58376 (2020) can be accessed under GSE150041).

