## Supplementary material for "Molecular determinants of cardiac lymphatic dysfunction in a chronic pressure-overload model": Suppl methods, Figures and Table S1

26th Feb 2025

#### List of Supplementary material

| | |
|---|---|
| Figure S1 | Marker genes and cell cycle phase distribution of cardiac endothelial cells [BALB/c] |
| Figure S2 | Distinguishing EC markers related to metabolic and MHC class I genes [BALB/c] |
| Figure S3 | EC cluster differences in key genes involved in vascular barrier regulation [BALB/c] |
| Figure S4 | Frequency of expression of LEC marker genes [BALB/c] |
| Figure S5 | Differentially-expressed genes post-TAC in cardiac BEC and vBEC clusters [BALB/c] |
| Figure S6 | Subpopulation analyses of cardiac BECs post-TAC [BALB/c] |
| Figure S7 | Cardiac LEC expression of “immune LEC” marker genes [BALB/c] |
| Figure S8 | Differentially-expressed genes in cardiac LECs post-TAC shared with IL1β-stimulated LECs |
| Figure S9 | Cardiac LEC expression profiles in AngII model [C57] |
| Video 1 | Lymphatic CCL21 expression in healthy hearts [BALB/c] |
| Video 2 | Lymphatic valves in healthy hearts [C57] |
| Table S1 | Mean transcripts and genes per cell [BALB/c; C57] |
| Table S2 | Marker genes of global LEC, BEC and vBEC clusters in healthy hearts [BALB/c] |
| Table S3 | Marker genes of BEC subpopulations in healthy hearts [BALB/c] |
| Table S4 | Marker genes of LEC subpopulations in healthy or TAC hearts [BALB/c] |
| Table S5 | Full list of DEGs *TAC vs sham* in global LEC cluster [BALB/c] |
| Table S6 | Full list of DEGs *TAC vs sham* in global BEC cluster [BALB/c] |
| Table S7 | Full list of DEGs *TAC vs sham* in vBEC cluster [BALB/c] |
| Table S8 | ORA analysis of cardiac LECs post-TAC [BALB/c] |
| Table S9 | Full list of DEGs *TAC vs sham* in BEC subpopulations [BALB/c] |
| Table S10 | Full list of DEGs *TAC vs sham* in LEC subpopulations [BALB/c] |
| Table S11 | Full list of DEGs *TAC vs sham* in cardiac EC subpopulations [C57] |
| Table S12 | Full list of DEGs induced by IL-1β in confluent LEC cultures [Human] |

**Supplementary methods**

| | |
|---|---|
| Table S13 | antibodies and reagents used in tissue sections |
| Table S14 | antibodies and reagents used for whole-mount staining |
| Table S15 | antibodies and reagents used for FACS in mouse |

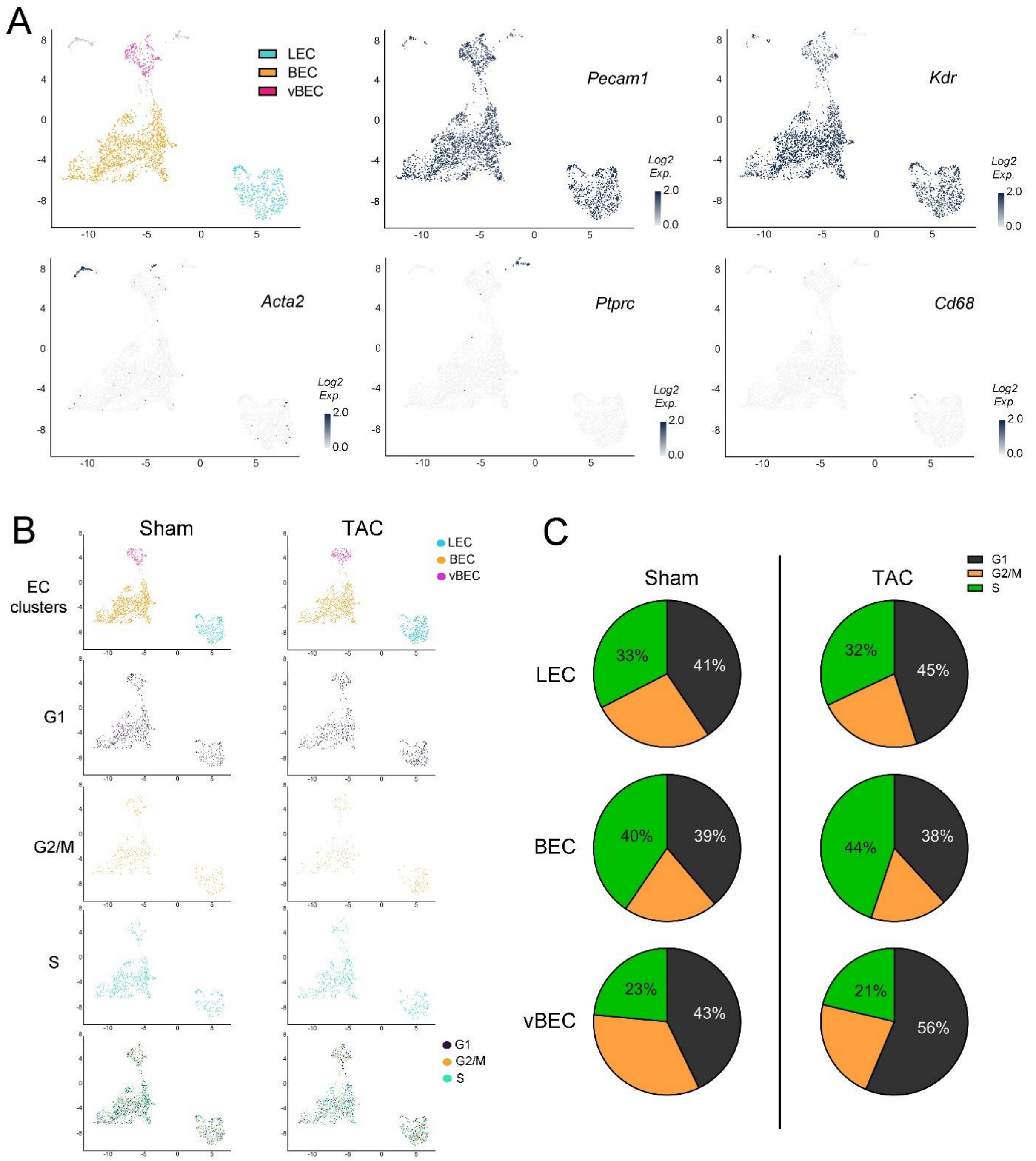

**Figure S1 Marker genes and cycle phase distribution of cardiac endothelial cells**

Visualization [UMAP, **a**] of cardiac EC clusters from healthy BALB/c mice. The three main EC clusters expressed vascular markers (*Pecam1, Kdr*), but not immune (*Ptprc, Cd68*) or mural cell marker (*Acta2*). Expression levels shown as *Log2 normalized read counts.* Visualization [UMAP, **b**] and quantification (**c**) of CellLoupe-based scoring of cell cycle phases in main EC clusters from healthy and post-TAC BALB/c hearts.

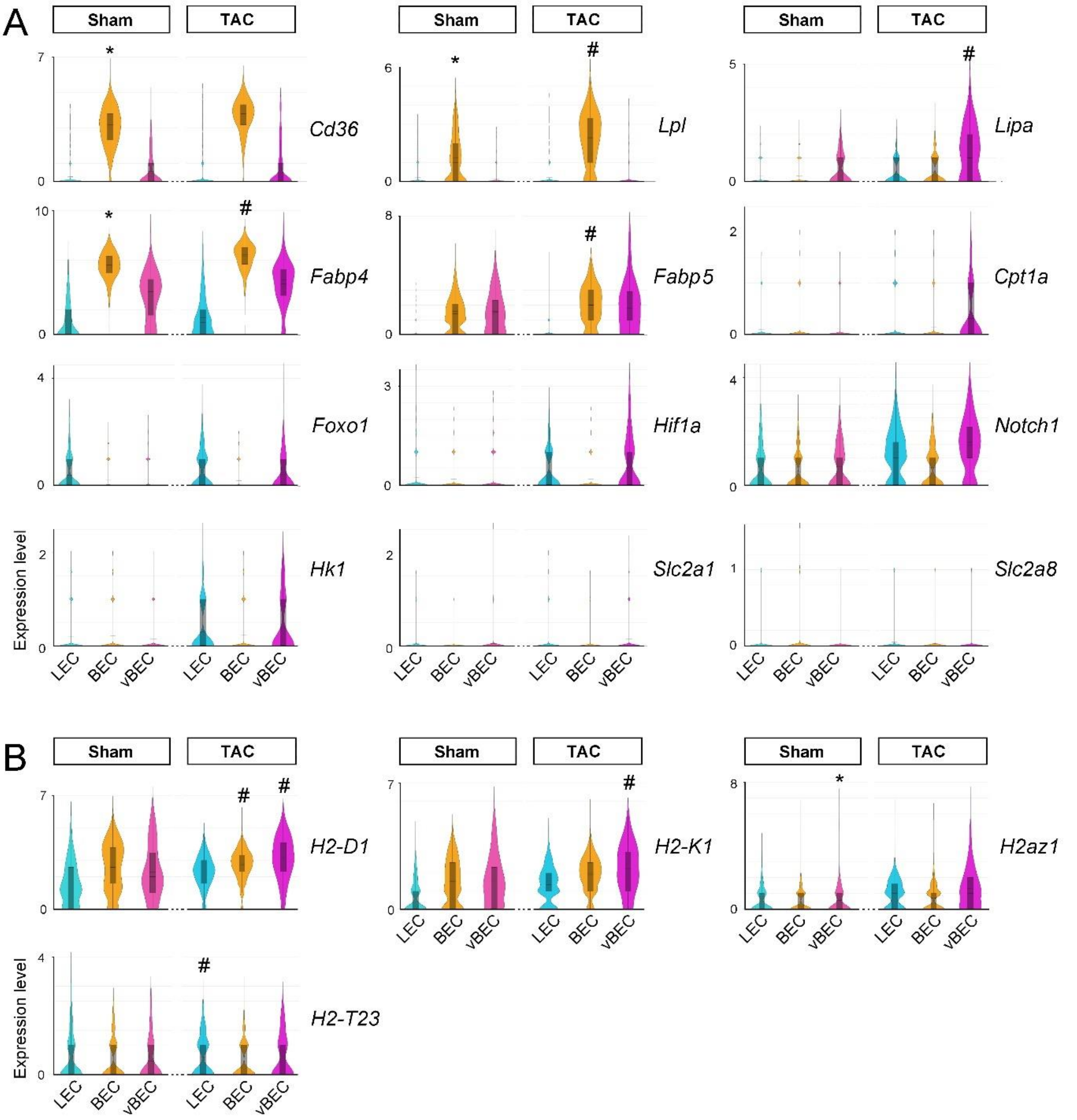

#### Figure S2 Distinguishing EC markers related to metabolic and MHC class I genes

Examples of expression levels [violin plot, *Log2 normalized read counts*] of genes involved in regulation of metabolism (**a**), either lipolysis (*Cd36, Lpl, Lpa, Fabp4, Fabp5, Cpt1a*) or glycolysis (*Foxo1, Hif1a, Notch, Hk1, Slc2a1, Slc2a8*), and main MHC class I molecules for antigen presentation (**b**) in cardiac EC clusters from healthy and post-TAC balb. Clusters significantly-enriched for a given gene are denoted (*****). For a full list of marker genes of EC clusters in healthy hearts, see **table S2**. Significantly altered genes post-TAC are denoted (#). For lists of DEGs post-TAC see **tables S5** (LEC), **S6** (BEC), and **S7** (vBEC). *Cpt1a*, Carnitine palmitoyltransferase*; Fabp,* Fatty acid binding protein; *Foxo1,* Forkhead box protein O; *Hif1a*, Hypoxia-inducible factor; *Hk1*, Hexokinase-1; *Lpl*, Lipoprotein Lipase; *Lpa*, Lipase A; *Slc2a1*, Glut1; *Slc2a8*, Glut8.

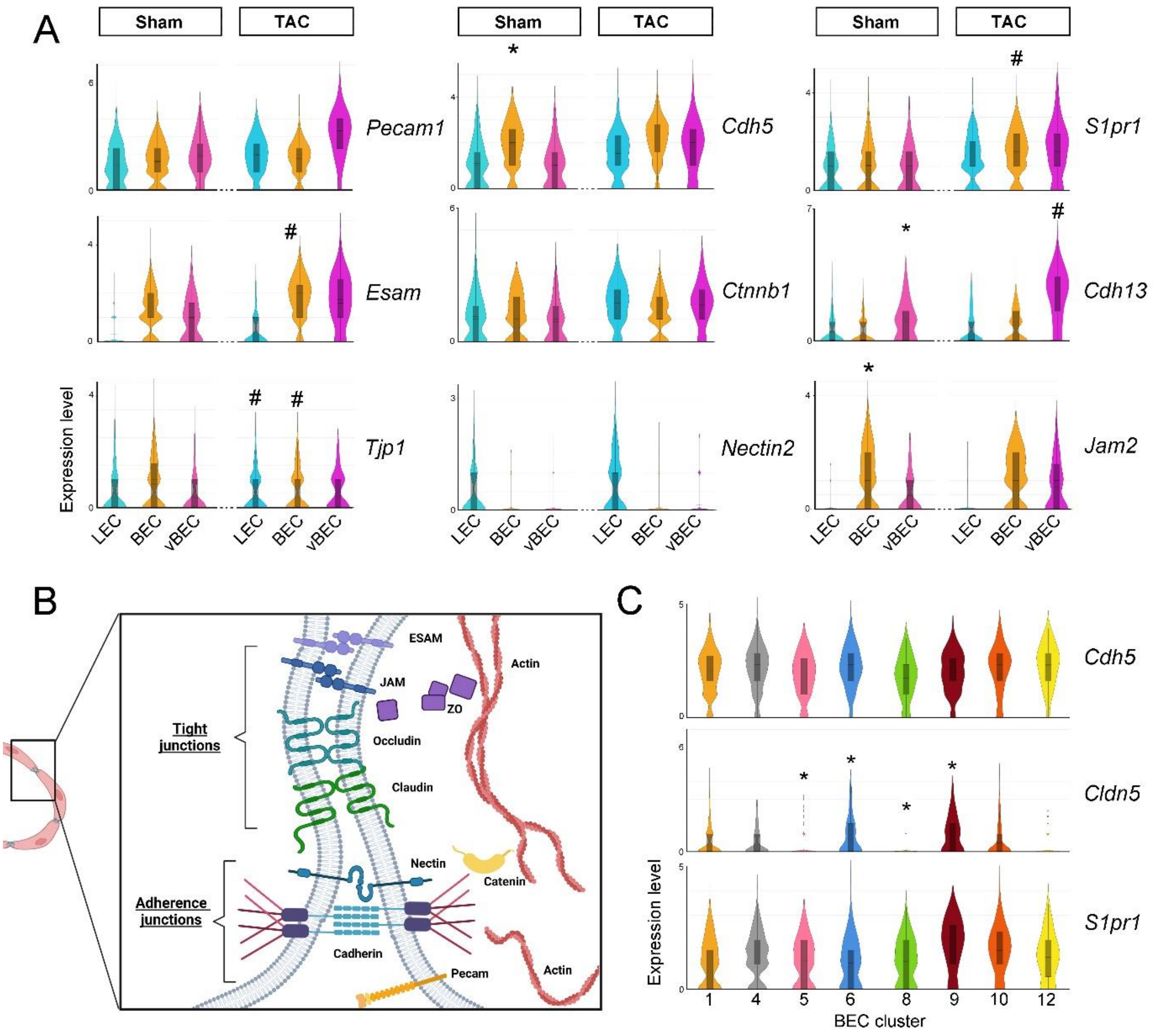

#### Figure S3 EC cluster differences in key genes involved in vascular barrier regulation

Examples of expression levels [violin plot, *Log2 normalized read counts*] of genes involved in adherence junctions and tight junctions in ECs from healthy and post-TAC balb mice (**a**). Clusters significantly-enriched for a given gene are denoted (*****); whereas genes significantly altered post-TAC are denoted (#). For a full list of EC marker genes, see **table S2**, and for DEGs post-TAC see **tables S5** (LEC), **S6** (BEC), and **S7** (vBEC). Schematic overview (created using BioRender) of junctional assembly (**b**). Examples of expression levels [violin plot] of barrier-relevant genes in healthy cardiac BEC subpopulations (**c**). For a list of BEC subpopulation marker genes see **table S3**, and for full list of DEGs in BEC subpopulations post-TAC, see **table S9**. *Cdh5*, VE-Cadherin; *Cdh13*, T-cadherin; *Ctnnb1*, β-Catenin; *Esam*, Endothelial cell adhesion molecule; *Jam*, Junctional adhesion molecule; *S1pr1*, Sphingosine-1-phosphate receptor 1; *Tjp1*, Tight junction protein-1/ZO-1.

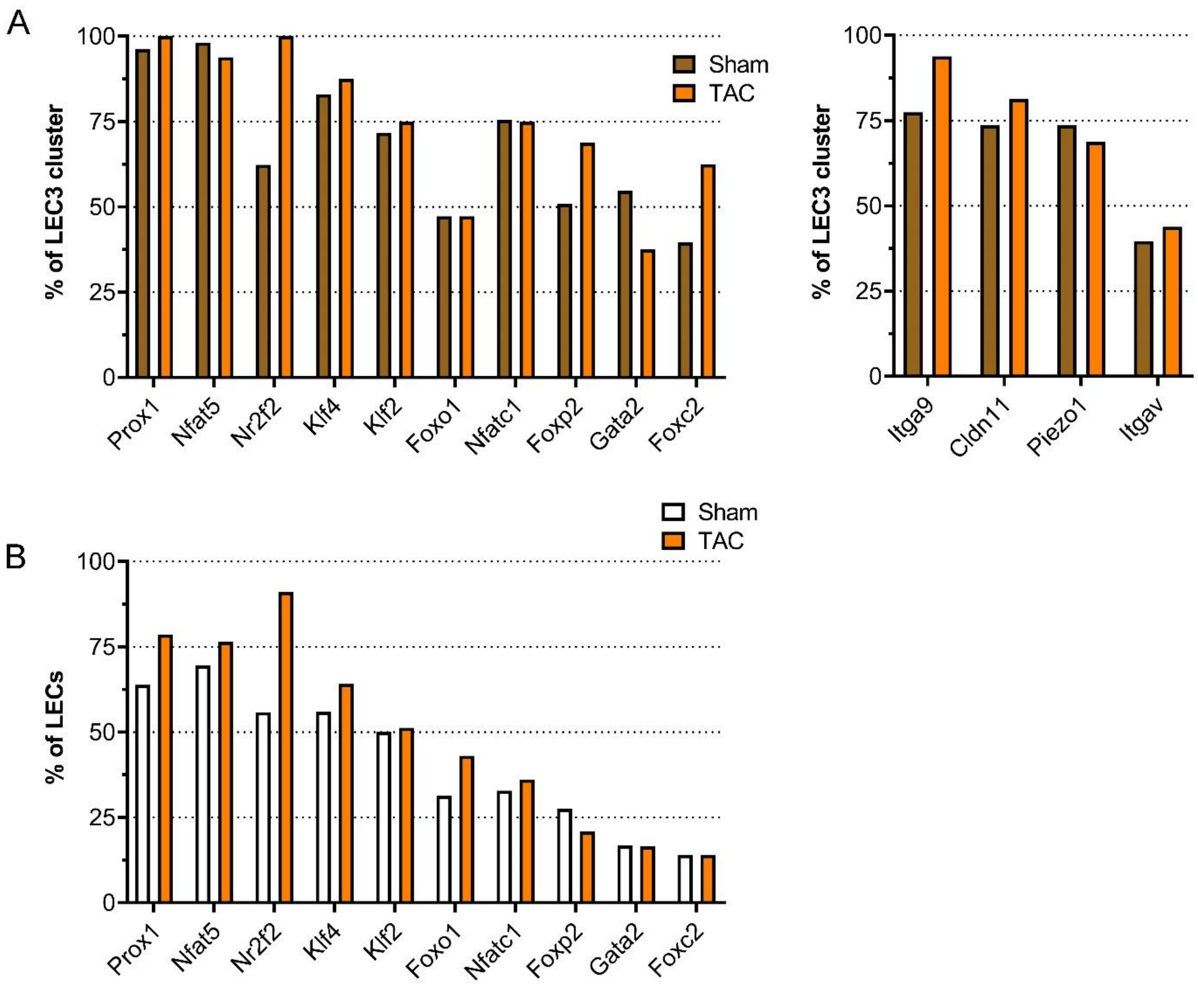

#### Figure S4 Frequency of expression of LEC marker genes

Examples of frequency of expression of lymphatic transcription factors and valvular marker genes in the cardiac LEC3 subpopulation (**a**) and in the global LEC cluster (**b**) in healthy and post-TAC BALB/c mice. For a full list of LEC marker genes, see **table S4**.

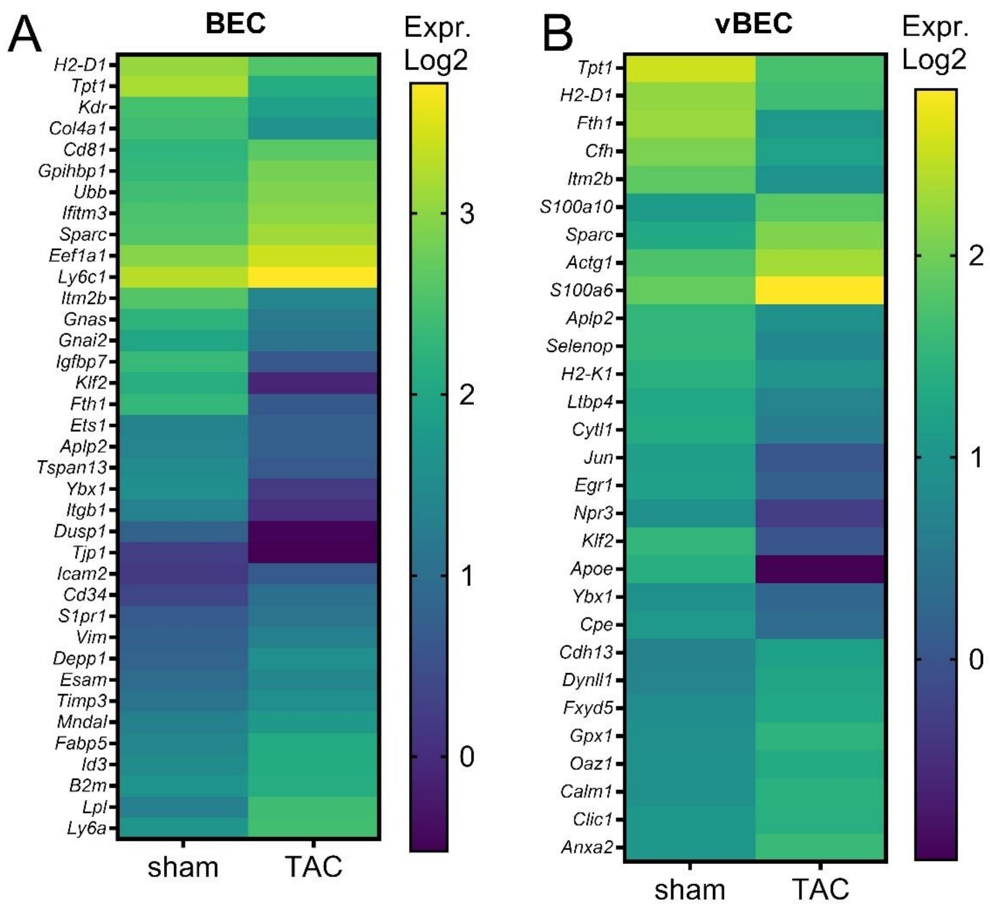

**Figure S5 Differentially expressed genes post-TAC in cardiac BEC and vBEC clusters**

Examples of genes differentially expressed post-TAC [heatmap, *Log2 normalized read counts*] in cardiac BECs (**a**) and vBECs (**b**), in BALB/c mice. For a full list of DEGs see **tables S6** and **S7**, respectively.

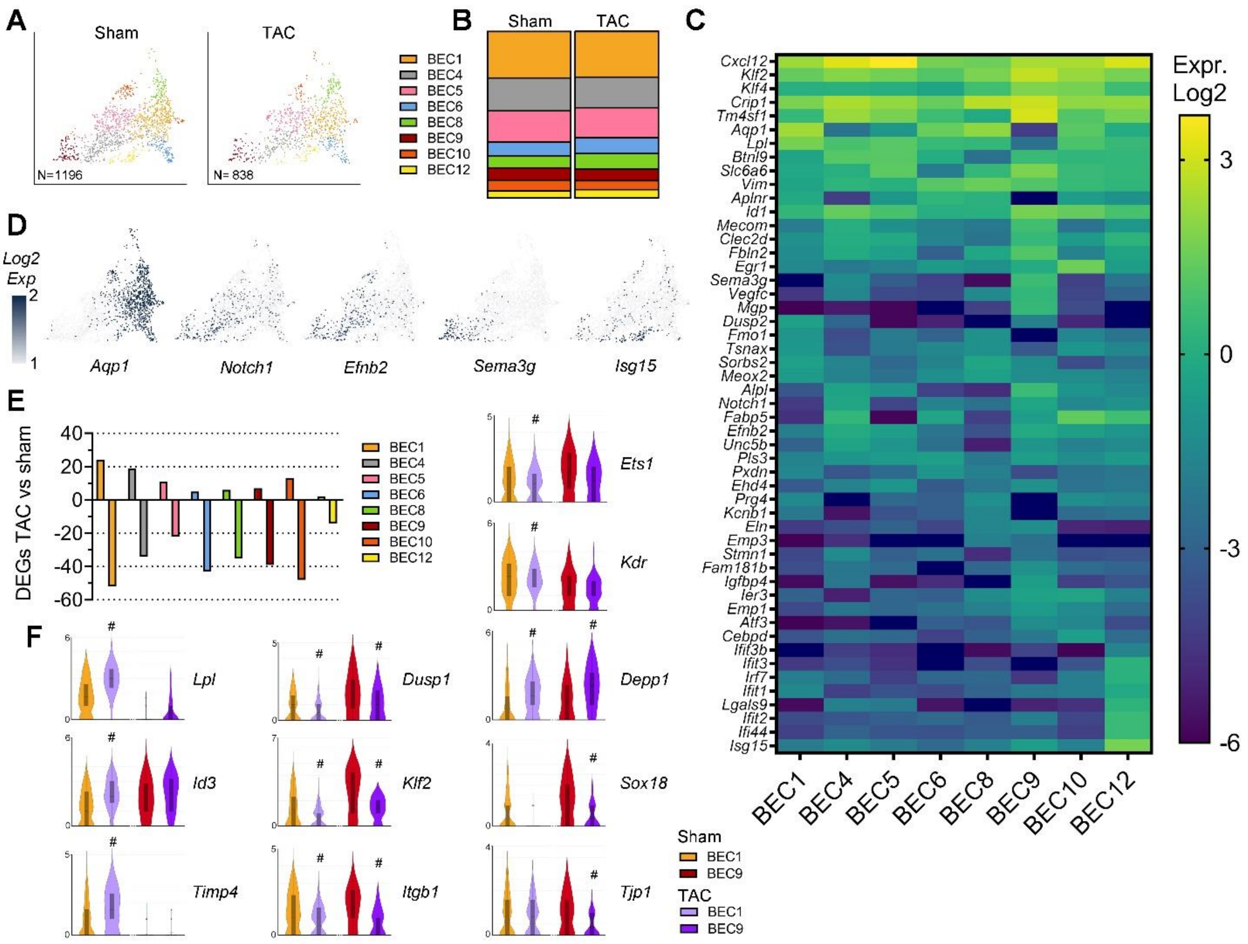

#### Figure S6 Subpopulation analyses of cardiac BECs post-TAC in Balb/c

Visualization [umap] of cardiac BEC subpopulations in healthy and post-TAC BALB/c mice (**a**). BECs from healthy and TAC-operated mice clustered into 8 subpopulations, with the majoritarian BEC1 cluster representing around 30% of cells in both sham and TAC hearts (**b**). Examples of gene expression levels [heatmap, *Log2 transformed normalized read counts*] of marker genes for BEC clusters (**c**). Capillary gene markers were enriched in clusters BEC1, 5, 6, 8, and 10, while arterial gene markers were enriched in clusters BEC4, 5, 9, and 10, and the rare BEC12 cluster displayed an interferon-type signature. For list of marker genes see **table S3**. Examples of distribution [umap] of marker genes (**d**). Quantification of DEGs identified post-TAC in each BEC cluster (**e**). Examples of gene expression levels [violin plots] in sham (*orange* or *red*) and TAC (*purple*) groups in BEC1 and BEC9 clusters (**f**). Significant genes indicated (#) for the respective cluster. For list of DEGs for BEC clusters see **table S9**.

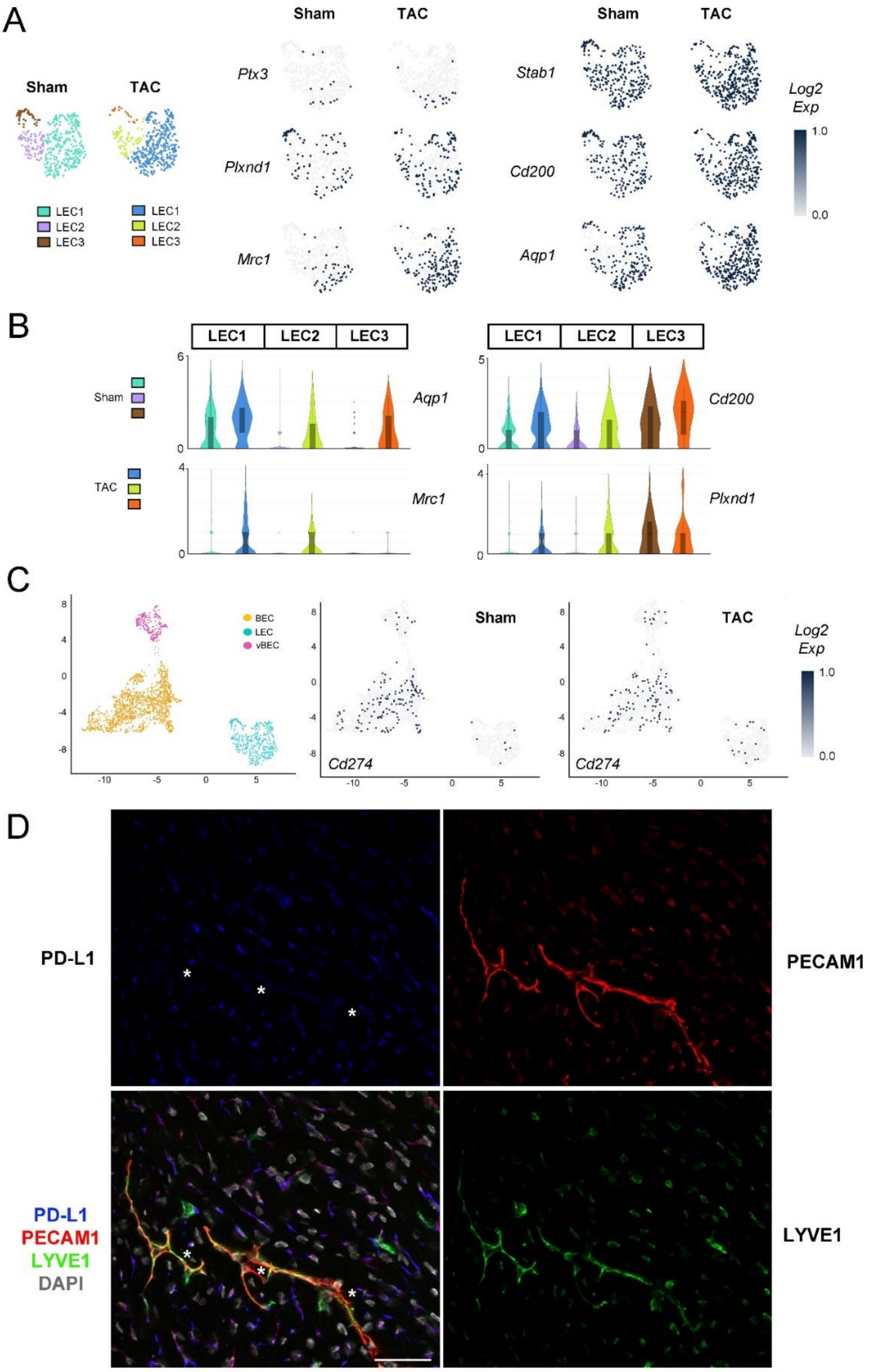

**Figure S7 Cardiac LEC expression of "immune LEC" marker genes**

Visualization [umap] of cardiac LEC subpopulations in healthy and post-TAC BALB/c mice, and LEC cluster distribution of marker genes proposed for immune" LEC (iLEC)[1] (**a**). Gene expression levels [violin plot, *Log2 normalized read counts*] of proposed markers analyzed in LEC clusters in healthy and post-TAC Balb/c mice (**b**). Cardiac EC expression distribution [umap] of Cd274 (Pdl1) in BECs, vBECs, and LECs in healthy and post-TAC Balb/c mice (**c**). Vascular PD-L1 protein levels (**d**) evaluated by immunohistochemistry in cardiac section at 8 weeks post-TAC in Balb/c. PD-L1, *blue*; Lyve1, *green*; CD31, *red*: DAPI, grey. Scale bar 50 µm. White Asterix points to lymphatics displaying low PD-L1 expression. *Bottom left panel*: Purple cells, CD31$^{+}$ blood vessel capillaries expressing Pd-L1; Green cells, Lyve1$^{+}$ CD31$^{neg}$ macrophages.

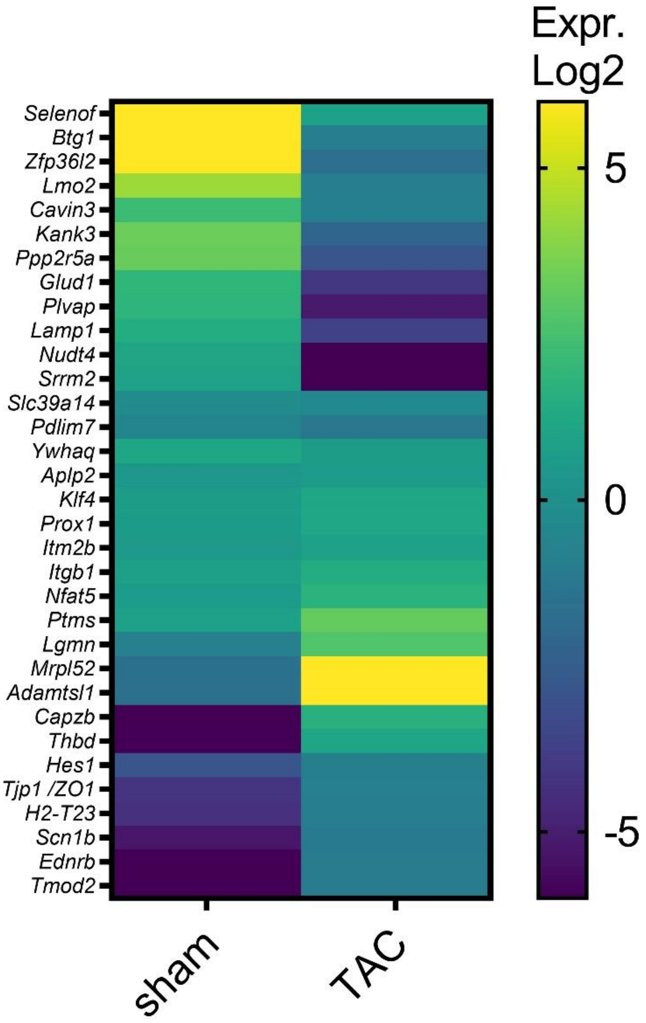

**Figure S8 Differentially-expressed genes in cardiac LECs post-TAC shared with IL1β-stimulated LECs**

Examples of expression levels [heatmap, *Log2 normalized read counts per cluster*] of genes significantly altered post-TAC in BALB/c mice. The selected genes were all similarly altered *in vitro* in IL1β-treated human LECs. For a list of all DEGs in human LECs, see suppl. **Table S12**.

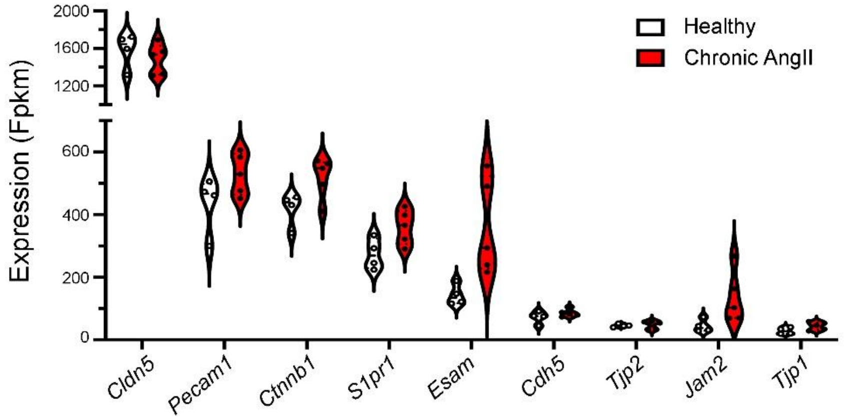

**Figure S9 Cardiac LEC expression profiles in AngII model**

Expression level changes [violin plot, *Fpkm counts*] in vascular barrier-related genes in cardiac LECs isolated for bulk RNAseq (n=4-5 samples/group) following pressure-overload induced by chronic AngII infusion for 6 weeks in C57 mice (adapted from *Song et al*.[2] GSE150041).

**Table S1 - Mean transcripts and genes per cell**

| **BALB/c ECs** | ***N* (cells)** | **Σ nUMI (reads)** | **Σ nGene** | **nUMI / nGene** |
|---|---|---|---|---|
| Healthy LEC | *402* | 3079 | 1400 | 2,2 |
| Healthy BEC | *1196* | 2514 | 1324 | 1,9 |
| Healthy vBEC | *216* | 3002 | 1323 | 2,3 |
| **Global healthy** | ***1814*** | **2699** | **1341** | **2,0** |
| TAC LEC | *453* | 5027 | 2075 | 2,4 |
| TAC BEC | *833* | 3006 | 1468 | 2,0 |
| TAC vBEC | *168* | 6811 | 2338 | 2,9 |
| **Global TAC** | ***1459*** | **4096** | **1761** | **2,3** |
| **C57 ECs** | *N* (cells) | Σ nUMI (reads) | Σ nGene | nUMI / nGene |
| Healthy LEC | *150* | 4023 | 1837 | 2,2 |
| Healthy BEC | *380* | 3008 | 1470 | 2,0 |
| Healthy vBEC | *104* | 5018 | 2101 | 2,4 |
| **Global healthy** | ***634*** | **3564** | **1656** | **2,2** |
| TAC LEC | *81* | 7606 | 2693 | 2,8 |
| TAC BEC | *704* | 4712 | 1992 | 2,4 |
| TAC vBEC | *107* | 8006 | 2733 | 2,9 |
| **Global TAC** | ***892*** | **5370** | **2145** | **2,5** |

Average Read counts and Features per cell in global EC population or main cluster

### Supplementary methods

#### Experimental Model

Female BALB/c and C57BL6/J mice (22-24 g) were obtained from Janvier. Animal housing and experiments were in accordance with European Directive 2010/63/EU on the protection of animals, and the study was approved by the Normandy University ethical review board Cenomexa according to French and EU legislation (APAFIS #**23175**-2019112214599474 v6; APAFIS #**32433**-2022070712508369 v2). Minimally-invasive transversal aortic banding constriction (TAC) was performed on 8-week-old mice, as previously described.[3] Mice were anaesthetized by intraperitoneal injection of ketamine (100 mg/kg Imalgene®) and xylazine (10 mg/kg Rompun® 2%, Bayer Health Care) and placed on mechanical ventilation. The operator performed a minimal thoracotomy with an incision at the level of the first intercostal space. The aortic arch was visualized under low-power magnification. A snare, made of 7-0 polypropylene suture, was passed under the aorta between the origin of the right innominate and left common carotid arteries. Two suture bands were placed side-by-side to create an elongated stenosis and prevent internalization of the suture, as described.[4] A bent 26-gauge needle was placed next to the aortic arch, and the sutures were snugly tied around the needle and the aorta. After banding, the needle was quickly removed. The skin was closed, and mice recovered on a warming pad until fully awake. The sham procedure was identical except that the aorta was not banded. Buprenorphine (50 µg/kg, Buprecare®, Axcience) was injected subcutaneously 6 hours after surgery and twice per day until 3 days post-operation. Euthanasia was performed by pentobarbital overdose (100 mg/kg Euthoxin).

#### Immunohistochemistry and epifluorescence microscopy

Murine cardiac samples were sectioned into a central slice, which was snap-frozen. Cardiac sections were cut on a cryostat (8 µm thickness) and collected on SuperFrost plus glass slides. After fixation in acetone for 10 min, non-specific binding sites were blocked in 3% BSA in PBS, followed by Biotin-Avidin Blocking kit (Thermo Scientific) when streptavidin (SA)-conjugates were used to detect biotinylated secondary antibodies. Primary antibodies (*see* **Table S13**), diluted in 1% BSA in PBS, were incubated on the sections at r.t. for 1h, followed by repeated washing in PBS and incubation with secondary antibodies for 30 minutes to 1h. Multi-stainings were performed sequentially, and negative controls included omission of primary antibodies. Slides were mounted in Vectashield containing DAPI, and images were acquired using x20 or x40 objectives on a Zeiss epifluorescence microscope (AxioImager J1) equipped with an apotome and Zen 2012 software (Zeiss), or using x63 objectives on a Leica Thunder Tissue 3D miroscope. Images were analyzed using Fiji imaging software (NIH) by an operator blinded to the groups.

**Table S13**–antibodies and reagents used in tissue sections

| antigen | article | supplier | species reactivity | host | dilution | Working conc (µg/mL) |
|---|---|---|---|---|---|---|
| **alpha SMA-FITC** | *F3777* | Sigma Aldrich | mouse | mouse | 1/100 | **28** |
| **CD31/PECAM** | *553371* | BD | mouse | rat | 1/100 | **0.6** |
| **CD68** | *14-0681-82* | eBioscience | mouse | rat | 1/800 | **5** |
| **CCL21** | *AF457* | RnD systems | mouse | goat | 1/100 | **5** |
| **Claudin-5** | *34-1600* | Invitrogen | mouse | rabbit | 1/400 | **0.6** |
| **F4/80** | *MCA497R* | Abd Serotec | mouse | rat | 1/200 | **5** |

| | | | | | | |
|---|---|---|---|---|---|---|
| **LYVE-1** | *103-PA50* | Reliatech | mouse | rabbit | 1/1000 | **0.4** |
| **Biotinylated LYVE-1** | ALY7 13-0443-82 | eBiosciences | mouse | rat | 1/500 | **0.5** |
| **CD206/MRC1** | *ab64693* | Abcam | mouse | rabbit | 1/2500 | **4** |
| **Pdl1** | 124302 | Biolegend | mouse | rat | 1/50 | **10** |
| **Podoplanin** | *14-5381-82* | eBioscience | mouse | hamster | 1/10 000 | **1** |
| **Reelin** | *AF3820* | RnD systems | mouse | goat | 1/50 | **4** |
| **VEGF-C** | *ab9546* | Abcam | mouse | rabbit | 1/500 | **2** |
| **VCAM-1** | *sc-19982* | SantaCruz | mouse | rat | 1/500 | **0.4** |
| **ICAM-1** | *14-0541-82* | eBioscience | mouse | rat | 1/100 | **5** |
| **VE-cadherin** | *AF1002* | RnD systems | mouse | goat | 1/200 | **1** |
| **WGA** | *FP-CE8070* | Interchim | | | 1/100 | **1** |

| **reactivity** | article | supplier | Fluorochrome | Working conc (µg/ mL) |
|---|---|---|---|---|
| **Donkey anti-Rat** | *712-545-153* | Jackson Immunoresearch | **AF488** | **3** |
| **Donkey anti-Rat** | *712-166-153* | Jackson Immunoresearch | **Cy3** | **3** |
| **Goat anti-Rat** | *A-21247* | Thermo Fisher Scientific/ Invitrogen | **AF647** | **0.8** |
| **Donkey anti-Rabbit** | *711-165-152* | Jackson Immunoresearch | **Cy3** | **3** |
| **Donkey anti-Rabbit** | *711-605-152* | Jackson Immunoresearch | **AF647** | **1.5** |
| **Donkey anti-Goat** | *A50-201D2* | Interchim | **DYLIGHT488** | **1.3** |
| **Donkey anti-Goat** | *A50-201D3* | Interchim | **DYLIGHT 550** | **1.3** |
| **Streptavidin** | *FP-CA5570* | Interchim | **Fluoprobe 547** | **0.7** |
| **Streptavidin** | *FP-CA5640* | Interchim | **Fluoprobe 647** | **0.7** |
| **Goat anti-Hamster** | *A-21110* | Thermo Fisher Scientific / Invitrogen | **AF488** | **0.8** |

#### Whole mount immunohistochemistry

Whole mount staining was performed, as previously described[3]. Briefly, prior to sacrifice, deeply anesthetized mice were perfused with warm saline solution, followed by perfusion-fixation with warm 3% paraformaldehyde (PFA). Hearts were removed and postfixed (3% PFA) for 6h. Following dehydration in graded methanol baths, and post-fixation in Dent's fixative, samples were bleached in graded H2O2 baths. After extensive blocking of nonspecific binding sites and tissue permeation with Triton-X100, cardiac lymphatics were visualized using antibodies (*see* **Table S14**) reactive against Lyve1, Popoplanin, CCL21, Podocalyxin and Reelin, followed by fluorescence-coupled secondary antibodies. Blood vasculature was visualized using anti-alpha-smooth muscle actin and Podocalyxin antibodies. Extensive washing was performed to remove nonspecific binding. Hearts were clarified using a modified iDISCO+ protocol, as described4, based on incubation in graded methanol baths followed by incubation in dichloromethane (DCM, Sigma-Aldrich, #270997-12X100ML) and dibenzyl ether (DBE, Sigma-Aldrich, #108014-1KG) before light sheet and confocal imaging.

**Table S14–**antibodies and reagents used for whole mount staining

| antigen | article | supplier | species reactivity | host | dilution | Conc (µg/mL) |
|---|---|---|---|---|---|---|
| **alpha SMA-Cy3** | *C6198* | Sigma Aldrich | mouse | mouse | 1/500 | **3** |
| **CCL21** | *AF457* | RnD systems | mouse | goat | 1/100 | **5** |
| **LYVE-1** | *103-PA50AG* | Reliatech | mouse | rabbit | 1/500 | **0.8** |
| **Biotinylated LYVE1** | ALY7 13-0443-82 | eBiosciences | mouse | rat | 1/500 | **0.5** |
| **Podocalyxin** | AF1556 | RnD systems | mouse | goat | 1/200 | **1** |
| **Podoplanin** | *14-5381-82* | eBioscience | mouse | hamster | 1/500 | **0.1** |
| **antigen** | **article** | **supplier** | Fluoro-chrome | | dilution | Conc (µg/ mL) |
| **Donkey anti-Rabbit** | *711-165-152* | Jackson Immunoresearch | **Cy3** | | 1/500 | **3** |
| **Donkey anti-Rabbit** | *711-605-152* | Jackson Immunoresearch | **AF647** | | 1/500 | **3** |
| **Donkey anti-Goat** | *A50-201D3* | Interchim | **DYLIGHT550** | | 1/500 | **3** |
| **Donkey anti goat** | *705-585-147* | Jackson Immunoresearch | **Cy3** | | 1/400 | **1.25** |
| **Streptavidin** | *FP-CA5570* | Interchim | **Fluoprobe547** | | 1/300 | **0.7** |
| **Goat anti-Hamster** | *A21113* | Thermo Fisher Scientific / Invitrogen | **Cy3** | | 1/1000 | **0.8** |

#### Light sheet and confocal microscopy

3D imaging was performed by light sheet microscopy as described[3]. Acquisitions were performed with an ultramicroscope II (LaVision BioTec) or Blaze (Miltenyi) using the ImspectorPro software (LaVision BioTec). The lightsheet was generated by a laser (wavelengths 561 or 640 nm, Coherent Sapphire Laser, LaVision BioTec) focused using two cylindrical lenses. A binocular stereomicroscope (MXV10, Olympus) with an *x*2 objective (MVPLAPO, Olympus) was used at different magnifications (*x*0.8 and *x*4). A dipping cap protective lens, including correction optics for MVPLAPO x2 objective, was applied for working distances inferior or equal to 5.7mm.

Samples were placed in an imaging reservoir made of 100% quartz (LaVision BioTec) filled with DBE and illuminated from the side by the laser light sheet. A PCO Edge SCMOS CCD camera (2560× 2160 pixel size, LaVision BioTec) was used to capture images. The step size between each image was fixed at 6 µm (*x*0.8 zoom) or 2 µm (*x*3.2 zoom). All tiff images were generated in 16-bit.

For confocal laser scanning microscopy, imaging was performed with an upright fixed-stage TCS SP8 confocal microscope (Leica Microsystems, France) equipped with multiple laser lines (wavelengths 561 or 640 nm). In order to acquire deep confocal views, a *x25* objective was used (numerical aperture 0.95, working distance 2500 µm, water immersion). Images were captured using a hydrid detector (Hamamatsu) in photon counting z-stack mode. Maximal intensity projection views were generated with Fiji Image J software (1.54g, Java 1.8.0_322, 64-bit).

Images, 3D volume, and movies from Lightsheet microscopy were generated using Imaris x64 software (version 8.0.1, Bitplane). Z stack images were first converted to imaris file (.ims) using ImarisFileConverter and 3D reconstruction was performed using the “volume rendering” function. To facilitate image processing, images were converted to 8-bit format. Optical slices were obtained using the “orthoslicer” tool. 3D pictures and movies were generated using the “snapshot” and “animation” tools. Movie reconstruction with .tiff series were performed with Fiji Image J software.

#### Fluorescence-activated cell sorting

At 8 weeks post-TAC, animals were euthanized by barbiturate over-dose (100 mg kg $^{-1}$ intraperitoneal, Eutoxin), and hearts were recovered after perfusion with warmed physiological saline through the abdominal aorta. LV samples were minced with scalpels and placed in ice-cold DMEM medium. Single cell suspensions were prepared, as described[3], by 30 min incubation in tissue-dissociating solution (125 U/mL collagenase type XI, *#C7657*, Sigma; 450 U/mL collagenase type I, *#C0130*, Sigma; 60 U/mL hyaluronidase type I, *#H3506*, Sigma; and 60 U/mL DNase 1 *#DN25*, Sigma) using gentleMACS Dissociator (MACS; Miltenyi Biotec, Auburn, CA). Digested tissues were washed with DMEM culture medium and filtered to remove undigested tissue pieces (80 µm mesh, and 40 µm mesh, BD Biosciences).

Cardiac single cell suspensions were prepared in FACS buffer (5% BSA in PBS). To block nonspecific binding of antibodies to Fcγ receptors, isolated cells were first incubated with Fc-Block for 15 minutes at 4°C. Subsequently, cells were stained with specific antibody cocktails (*see* **table S15**) for 20 minutes at r.t, followed by washing with FACS buffer and resuspension in 700 µL phenol red-free RPMI with 5% FCS for sorting.

FACS sorting was performed on an FACS ARIA II (BD Biosciences). BECs were defined as live $CD45^-/CD31^+/Lyve1^-$ cells, whereas LECs were identified as $CD45^-/CD31^+/Lyve1^+/Pdpn^+$ cells. 3000-100 000 cells were collected into 50-500 µL 100% FCS and kept on ice prior to scRNAseq. Cells were then centrifuged and resuspended in PBS at a concentration of 1000 cells/ µL for inclusion in 10X Genomics pipeline.

**Table S15**– antibodies and reagents used for FACS in mouse

| Antigen | Fluorochrome | host | Source | article |
|---|---|---|---|---|
| **CD16/32 (Fc block)** | - | rat | BD Pharmingen | 553142 |
| **Fixable Viability Dye (L/D)** | **eFluor™ 455UV** | - | ebioscience | 65-0868-14 |
| **CD31** | **APC** | rat | BD Pharmingen | 551262 |
| **CD45** | **PerCP** | rat | Sony Biotechnology | 1115650 |
| **Lyve1** | **PE** | rat | RnD systems | FAB212SP |
| **Podoplanin** | **A488** | hamster | Sony Biotechnology | 1320070 |
| **Podoplanin** | **A488** | hamster | Ozyme/Biolegend | 127406 |

#### scRNAseq: sample preparation, sequencing, and data analysis

A 1:3 mix of cardiac LECs and BECs were included in each sample prepared following the 10X Genomics pipeline. Briefly, we charged 16’000 cells per reaction. For each group, 1-2 reactions were performed using cells from 10 pooled mouse hearts. The 10X Genomics Chromium Next Gem Single Cell 3' library kit v.3.1 was used to prepare barcoded mRNA following the guidelines from the supplier. Single Index kit T Set A was used to label each reaction. Samples were loaded onto 3 separate Illumina kit High-throughput 2x75 (<400 million reads) flow-cells and sequenced using Illumina Nextseq, using the settings of: *Read1*: 28 cycles; *Read2*: 8 cycles; *Read3 (i7)*: 114 cycles.

Raw FASTQ files, integrating single index and cell barcodes, were generated and demultiplexed from BCL sequencing output files using bcl2fastq (v2.20.0)[5]. FastQC (v0.11.9)[6] was used for quality control, revealing >98% perfect index and high-quality reads (86% bases >= Q30), requiring no trimming or filtering, with the exception of BALB/c sham1 sample trimmed using FASTP (v0.20.0)[7] to remove adapters, poly-A and poly-G present in Reads 2. In each sample, the STARsolo RNAseq aligner (v2.7)[8] mapped over 88% of reads to the mouse reference genome (GRCm39.111, Ensembl ftp server), generating a feature-barcode matrix (including *Read 1*, *Read 2,* and *i7* index) for each sample, based on feature (gene) counting per cell.

Data filtering, normalization, integration, clustering, and differential analysis were performed using the package Seurat (version 5.0.2)[9] in R (version 4.1.2). For low-quality data filtering, genes expressed by <10 cells, and cells expressing <500 genes for Balb/c samples or <300 for C57 samples, were excluded, as were cells with UMI numbers <500 or cells with >10% mitochondria-derived UMI counts. Matrices were normalized using 'NormalizeData()' (LogNormalize, scale factor 10,000). The top 3000 highly-variable features were selected using 'FindVariableFeatures()' (vst method). To correct batch effects and integrate datasets, Seurat's anchor-based method with SNN clustering and Harmony[10] were applied. Cell cycle heterogeneity was mitigated with 'CellCycleScoring()' and mitochondrial UMI counts via 'SCTransform()'. The top 3000 variable genes were selected ('SelectIntegrationFeatures()'), and data integration was performed using 'FindIntegrationAnchors()' and 'IntegrateData()', producing integrated matrices.

Principal component analysis (PCA) was applied to the normalized and integrated data matrices, followed by Harmony to correct for any remaining batch effects. Principal components (PCs) were selected by accumulating their explained variance until reaching 90%, ensuring no unique component exceeded 5%. The first PC collection meeting these criteria was selected for downstream analysis. This included 42 dimensions for Balb/c data and 43 dimensions for C57 data. Unsupervised cell clustering was performed using 'FindNeighbors()' and 'FindClusters()', applying the Leiden algorithm[11,12]. Cell clusters were visualized using 2D UMAP (Uniform Manifold Approximation and Projection). The resolution was increased from 0.1 to 1.0 in increments of 0.1. The optimal resolution was determined using the clustree functions from the R clustree package[13], i.e., 0.6 for Balb/c data, and 0.9 for C57 data. Cell type automatic annotation was performed using the SingleR and Celldex R packages[14] with the ImmGen reference[15], before manual validation by inspecting key marker genes for each cluster. The filtered, integrated, normalized transcriptomes clustered into 11 and 9 main populations, respectively. Among 3908 total sequenced cells from Balb/c mice, 3436 cells remained post-filtering, including 1932 cells from sham-operated controls and 1504 cardiac cells from TAC-operated mice. Among 4178 total sequenced cells from C57 mice, 1577 cells remained post-filtering, including 665 cells from sham-operated controls and 912 cardiac cells from TAC mice.

After identification of the three main endothelial cell (EC) cluster, BEC and LEC clusters were subdivided into sub-clusters for a more detailed analysis of subpopulations. Cardiac EC marker gene identification was performed using 'FindMarkers' (test.use='MAST', logfc.threshold=0)[16]. Functional enrichment analysis was performed using the R package "*clusterProfiler"* (v4.10.1)[17] with over-representation analysis (ORA) method for Gene Ontology and Pathways (KEGG, Reactome). Statistical output from the processed dataset, including Marker genes, DEGs, and functional enrichment analyses, are provided in Supplementary **tables S2-S11**. DEGs were defined as genes up- or down-regulated by at least 0.2 log2 fold-change and a *padj*. <0.05. Only DEGs whose mean normalized read counts surpassed 0.3 are included.

Processed data (*expression_data_value.csv* for BALB/c and *expression_data.csv* for C57) was exported for visualisation in CellLoupe (Loupe Browser 8.1.2, *10x Genomics*) used to generate UMAP, Heatmaps, Violin plots, and Volcano plots. At an estimated sequencing depth of ≈20 000 reads per cell, we detected 2500-6800 transcripts (unique molecular identifier, UMIs) and 1300-2300 genes (features) per cell in the processed datasets (**Table S1**). The scRNAseq dataset is available in the Gene Expression Omnibus repository (accession number: GSE290576 *Effect of pressure-overload on cardiac endothelial cells*).

#### *In vitro* cell culture

Human LECs (PromoCell, HDLEC-c, C-12216) were seeded in 6-well plates ($3\times10^5$/well) and grown to confluence in complete Endothelial Cell Basal Medium-2 (PromoCell, ECGM-MV2, CC-22121), including 5% FCS with VEGF, IGF, EGF, and FGF supplement, according to recommendations of the supplier. Next, cells were serum-starved (1% FCS, ECM w/o VEGF, ECGF, EGF) overnight, and exposed to 20 ng/mL recombinant human IL1β (200-01B, PeproTech) in 2 mL per well of serum-starvation media. After 24h cells were recovered and RNA extracted (Single Cell RNA Purification Kit, Norgen, 51800) for subsequent mRNAseq analyses and bioinformatics analyses by Biomarker Technologies (BMK) company (Munster, GE) for identification of differentially expressed genes. Briefly, poly-A mRNA-seq, sequencing length of 150 nt paired-end (PE150), were performed with IlluminaSeq. Reads were aligned to the human genome (GRCh38). Transcript quantification was performed with *featureCounts*[18], and differential analysis, including normalization of the original *readcount*, was performed with DESeq2 bioconductor package[19]. Statistical test (DESeq2 Wald test) was conducted on the expression matrix after standardization. Genes significantly up- or downregulated upon treatment were identified (for RNAseq data see **Suppl Table S12**), and results are reported as *Fpkm* and as log2 fold of control condition (1% FCS in ECM without cytokines and growth factors).
